# Distinct calcium, sodium, and water programs characterize mouse and human connecting tubule

**DOI:** 10.64898/2026.11.13.766571

**Authors:** Jessica P. Bahena-Lopez, Jeremiah V. Reyes, Mingzhen Su, Xiao-Tong Su, Joshua N. Curry, Pat Thaitongsuk, Sebastian Bachmann, Chao-Ling Yang, Jonathan W. Nelson, David H. Ellison

## Abstract

The kidney connecting tubule lies between the distal convoluted tubule and collecting duct. Even though short, it plays a key role in regulating calcium, sodium, and potassium excretion and water balance in health and disease. In rodents and humans, transitions between distal segments are gradual, making the assignment of functional properties to cell types difficult. To address this, we enriched mouse kidney distal tubule cells prior to performing single nucleus RNA-sequencing. Connecting tubule cells separated into three clusters, one expressed genes that are relevant to calcium transport, while another expressed genes that are relevant to water transport. The third was intermediate between the other two. While genes for electrogenic sodium transport were expressed throughout the connecting tubule, Hsd11b2, the gene responsible for aldosterone-responsiveness, was expressed primarily by the distal cluster. Immunocytochemistry confirmed the separation of calcium and water transport pathways and spatial transcriptomics confirmed the linear organization of this segment. Human transcriptomic data showed similar gene separation. The results show that rodent and human connecting tubules separate calcium and water transport, thereby limiting tubule fluid calcium accumulation. They also restrict aldosterone action to segments that are responsive to arginine vasopressin to maintain sodium homeostasis during antidiuresis.

## Introduction

The kidney connecting tubule (CNT) was identified by anatomists working before and during the first part of the last century as a unique kidney tubule segment interposed between the distal convoluted tubule (DCT) and the collecting duct.^1^ First called a ‘connecting segment’ (*verbindunsstück*), it was much later recognized as a functionally distinct subdivision of the distal tubule ^2^. Using isolated perfused rabbit tubules, Imai and colleagues found that the connecting tubule (CNT) exhibits electrophysiological and pharmacological properties that are different from both the DCT and the cortical collecting duct (CCD).^2^ In experiments primarily using rodents, other groups showed that distinct distal tubule segments responded differently to treatment with hormones.^3–7^

A complicating factor in these analyses was the structural and functional differences between species; in rabbit kidneys, used originally for perfusion ex vivo, the junctions between DCT, CNT, and CD were abrupt. In contrast, in rodents and humans, transitions between distal segments were gradual;^8^ further, in deep cortical nephrons, CNTs formed into arcades that joined multiple tubules, rather than being interposed between a single DCT and a collecting duct. As currently envisaged, the CNT is characterized as comprising connecting tubule cells and intercalated cells (IC), with the latter being important sites of acid and chloride movement.^9^ Yet the early structural analyses suggest that the segment is more mosaic, including at least four cell types.^1^

Early studies emphasized the special role that the CNT plays in reabsorbing calcium, with several calcium-regulating hormones active therein^5,6,10,11^ and ex vivo functional studies providing clear functional support.^12–14^ Although initial studies focused on the CNT, most contemporary reviews ascribe distal calcium transport more generally to the *distal convolution* in rodents and humans, a segment comprising both the DCT and CNT.^15,16^

A second function ascribed to the CNT is electrogenic sodium absorption and potassium secretion.^17^ While work using isolated perfused tubules originally pointed to the central importance of the cortical collecting duct in sodium and potassium transport, more recent work suggests that the CNT is the site of highest ENaC mediated sodium absorption and potassium secretion, at least under most conditions.^17,18^ Korbmacher and colleagues found that sodium transport along the proximal portion was constitutive, whereas it is aldosterone-sensitive more distally. Yang and colleagues also described two segments of CNT and introduced the terms CNTe (early CNT) and CNTas (aldosterone-sensitive CNT).^17^

A third role for the CNT is vasopressin-regulated water transport, but this function may be species-specific. Early work using rabbit CNT segments found low water permeability that was unaffected by arginine vasopressin (AVP).^2^ In contrast, studies using mouse and rat kidneys suggested that at least CNT cells do express aquaporin-2 (Aqp2). Wade and colleagues showed that some CNT segments express Aqp2, while others do not^19,20^ and observed that Aqp2 is also present along arcades.^21^ This group later named the segments according to aquaporin-2 expression as CNT1 and CNT2, with only the latter expressing the water channel.^22^ Most recently, transcriptomic analyses, first using dissected tubule segments, later using single cell and single nucleus RNA-Sequencing (snRNA-seq), and finally lineage tracing, has allowed investigators to revisit cell types along the distal tubule. Zhang and colleagues also found evidence for two segments of the CNT, one of which expresses Aqp2 and the other does not and used the terms CNT1 and CNT2.^23^

Thus, the CNT is believed to play important roles in sodium and calcium absorption, in potassium secretion, and in vasopressin-mediated water reabsorption, but these properties have been variably ascribed to subsegments and to overlapping and discrepant classifications. Here, we integrate findings from complementary approaches to clarify the structure of the rodent and human CNT and provide insight into the segregation of functional properties. Because the CNT comprises only a small minority of cortical cells, we combined dissected tubule staining, immunofluorescence of kidney sections, whole-kidney and distal tubule-enriched snRNA-Seq, and spatial transcriptomics to provide a comprehensive classification of CNT structure and postulate its functional implications. The results provide insights into aspects of CNT function and resolve critical gaps in that classification of CNT cell types.

## Methods

### Animals

All animal studies were approved by the IACUC of the Oregon Health and Science University (Protocol IP00286). Male C57BL/6 mice (10-12 weeks old) were housed in metabolic cages under a 12:12-hour light/dark cycle. To profile CNT cells Calbidin-Cre mice (B6;129S-Calb1tm2.1(cre)Hze/J; JAX stock no. 028532) were crossed with the INTACT line (B6;129-Gt(ROSA)26Sortm5(CAG-Sun1/sfGFP)Nat/J, JAX 021039, J).^24–26^

### Enriched snRNA-seq

Calb1-Cre mice were crossed with the INTACT reporter line to generate Calb1-Cre-INTACT mice with Cre-dependent GFP labeling of the nuclear envelope. Two male mice, aged 8 and 14 weeks, were used and had ad libitum access to water and a gel diet (Envigo TD170002) containing 0.08% magnesium. Kidneys were perfused with PBS to remove residual blood, and half of each kidney was snap-frozen for subsequent snRNA-seq. Nuclei were isolated from frozen tissue using the Chromium Nuclei Isolation Kit (10x Genomics; catalog no. 1000494) according to the manufacturer’s instructions. GFP-positive nuclei were enriched by fluorescence-activated nuclei sorting (FANS), as previously described su nelson ^27^. Briefly, isolated nuclei were stained with Vybrant DyeCycle Ruby for 15 minutes on ice, and Ruby-positive, GFP-positive nuclei were selected. A total of 200,000 nuclei were collected per sample, centrifuged at 500 × g for 5 minutes at 4°C, and resuspended for counting. Library preparation and sequencing were performed using the 10x Genomics platform as previously described ^27^. Sequencing data were processed using Cell Ranger (10x Genomics) to generate gene-by-nucleus count matrices for downstream analysis.

#### Data processing and clustering

snRNA-seq data were processed in R v4.3.2 using Seurat v5.0.1. Ambient RNA was corrected using SoupX v1.6.2. Genes detected in ≥3 nuclei and nuclei with ≥200 detected genes were retained; nuclei with ≥10,000 genes or ≥30,000 UMIs were excluded. After normalization, variable-feature selection, scaling, and principal component analysis, doublets were identified using DoubletFinder v2.0.4 with optimized pK and homotypic-adjusted expected doublet numbers. Singlets were retained, mitochondrial genes were removed, and PCA was repeated. Following quality control and doublet removal, samples were merged and integrated using canonical correlation analysis (CCA) in Seurat. The first 30 CCA-integrated dimensions were used for nearest-neighbor graph construction, graph-based clustering, and UMAP visualization. Cell types were annotated using canonical markers, as previously described by Chen et al. and Su et al.^9,27^ **Differential gene expression**. Cell-type-enriched genes were identified using Seurat’s FindAllMarkers function with the Wilcoxon rank-sum test, comparing each annotated cell type with all remaining nuclei. Positive markers were retained with detection in ≥25% of nuclei in either group, log2 fold change ≥0.5, and Bonferroni-adjusted P < 0.05. **Data visualization and module scores**. Figures were generated in R v4.5.2 using Seurat v5.5.1 and ggplot2 v4.0.3. Cell identities and gene expression were visualized using UMAP, feature plots, and dot plots. Dot size represented the percentage of expressing nuclei, and color indicated scaled average expression. Cell-type proportions were calculated relative to all annotated nuclei. Calcium, sodium, and water transport module scores were calculated using AddModuleScore and displayed on UMAP embeddings. Figures were exported as SVG or TIFF files.

#### Trajectory analysis

Pseudotime was inferred using Monocle3 v1.4.27 for DCT2, CNT1, CNT_T_, CNT2, and PC-CD nuclei, retaining the Seurat UMAP coordinates. A principal graph was learned without partitioning and rooted at the node most frequently associated with DCT2 nuclei. For heatmaps, nuclei with finite pseudotime were ordered accordingly; log-normalized gene expression was smoothed across ordered nucleus positions using splines with three degrees of freedom and standardized to gene-wise z-scores. Heatmaps were generated using ComplexHeatmap v2.26.1 without row or column clustering. **Transcriptional Program Identification.** To identify gene expression programs within the CNT independently of clustering, consensus non-negative matrix factorization (cNMF) was applied to subsetted CNT nuclei using raw counts for all detected genes (22,237 genes). Factorization followed the standard cNMF workflow ^28^ and K = 5 was selected. Three of the five identified programs corresponded to CNT states and were designated CNT1-like, CNT_T_-like and CNT2-like according to the state with the highest usage.

### Immunostaining kidney sections and isolated tubules

1Wild-type C57BL/6 and Calb1-Cre-INTACT mice were used for immunofluorescence studies. Kidney tissue and isolated kidney tubules were prepared and stained as previously described.^27^ Briefly, kidneys were perfusion-fixed with 3% paraformaldehyde in PBS (pH 7.4), cryoprotected overnight in sucrose/PBS (800 mOsm), and embedded in optimal cutting temperature compound. For tubule isolation, kidneys were perfused with L-15 medium containing type II collagenase (1 U/mL) and digested in the same medium for 30 minutes at 37°C. Tubules were isolated by microdissection, transferred onto poly-D-lysine–coated coverslips, and fixed with formalin for 30 minutes on ice. Kidney cryosections (5 μm) and isolated tubules were permeabilized with 0.5% Triton X-100 in PBS for 30 minutes and blocked with 5% milk in PBS for 30 minutes. Samples were incubated with primary antibodies diluted in blocking buffer for 1 hour at room temperature or overnight at 4°C. After three PBS washes, fluorophore-conjugated secondary antibodies were applied for 1 hour at room temperature. Samples were washed, counterstained with DAPI, and mounted using ProLong Diamond Antifade Mountant (Thermo Fisher Scientific). Images were acquired using a KEYENCE BZ-X800 microscope and a Andor BC43 spinning disk confocal microscope. Antibody sources, catalog numbers, and working dilutions are provided in Supplemental Table 1.

### Spatial transcriptomics

Publicly available Xenium spatial transcriptomic data from two mouse kidney samples were obtained from Bahena-Lopez et al.^29^ Data processing and visualization followed the previously described workflow, adapted for analysis of distal tubule populations.^29^ Briefly, distal convoluted tubule (DCT), connecting tubule (CNT), CD-principal cells (PC), IC, and urothelial cells were subsetted to examine populations represented in the Calb1-Cre–INTACT snRNA-seq dataset. Selected cells were re-clustered and annotated using marker genes defining the Calb1-Cre–INTACT populations, restricted to genes included in the Xenium panel. These included *Slc12a3*, *Egf*, *Calb1*, *Slc8a1*, *Trpv5*, *Aqp2*, *Aqp3*, and *Aqp4*, together with *Slc4a1*, *Slc2ca4*, and *Upk1b*. Attention was given to the relative expression of *Calb1*, *Slc8a1, Trpv5* and *Aqp2* when distinguishing DCT2, CNT, and principal cell populations. Spatial distributions and gene expression patterns were visualized using code adapted from the original study. Briefly, annotated populations were visualized on UMAP embeddings using Seurat’s DimPlot function. Marker expression was summarized using DotPlot, with dot size representing the percentage of cells expressing each gene and color indicating scaled average expression. Spatial maps were generated from Xenium cell segmentation polygons linked to cell annotations through their cell identifiers, using Seurat, sf, and ggplot2 v3.5.1. Whole-section and magnified views were used to display the anatomical distribution and local organization of the annotated populations. **CNT Cell type Concordance.** Correspondence between Xenium and Calb1-Cre–INTACT CNT clusters were assessed at both gene-and program-level. For gene-level concordance, we considered only well-detected reference genes defined as those present in the Xenium panel and detected (≥ 1 count) in at least 10% of Xenium CNT cells in both replicates (1,087 genes). Pairwise differentially expressed genes were identified in the snRNA-seq reference for each contrast (CNT1 vs CNT_T_, CNT_T_ vs CNT2 and CNT1 vs CNT2; Wilcoxon test, min.pct 0.10, |log2FC| ≥ 0.25, adjusted P < 0.05), yielding 182, 175 and 324 genes, respectively. For each contrast and replicate, we computed log2 fold changes in Xenium as in Seurat (log2 of mean counts per 10,000 + 1) and compared them with the reference by Spearman correlation, the fraction of genes changing in the same direction, and the slope of a linear regression of Xenium versus reference fold changes. For program-level concordance with Xenium, the same cNMF analysis was repeated on the same reference CNT nuclei, but restricted to genes on the Xenium panel (4,472 genes with non-zero counts). K = 6 was selected, and the resulting gene spectra were projected onto Xenium CNT cells with starCAT.^30^ **Cell-type proportions and anatomical distribution.** The distribution of each population across the cortex, outer and inner stripes of the outer medulla, and inner medulla was calculated as a percentage of the total number of cells belonging to that population in each sample. Percentages were summarized as the mean and standard deviation across the two biological replicates. **Spatial neighborhood analysis.** Neighboring cells were identified from their spatial centroid coordinates using a 20-μm radius. Self-neighbors and relationships between cells of the same type were excluded. For each source population, neighbor counts were normalized to the total number of heterotypic neighbors. Analyses were performed separately for all annotated distal populations across the tissue and for cortical DCT2, CNT1, CNT_T_, CNT2, and PC-CD cells. Sample-specific matrices were averaged with equal weight and displayed as heatmaps. **Trajectory analysis.** Cortical DCT2, CNT1, CNT_T_, CNT2, and PC-CD cells from both samples were combined using genes shared between the Xenium panels and analyzed with Monocle3 v1.3.7. Counts were log-normalized, reduced using 30 principal components, and embedded using UMAP. A principal graph was learned, and pseudotime was rooted at the graph node most frequently associated with DCT2 cells. Expression profiles for selected calcium, sodium, and water transport genes were smoothed separately for each sample using smoothing splines with three degrees of freedom over a shared pseudotime range. Profiles were averaged with equal sample weight, standardized to gene-wise z-scores, and displayed using ComplexHeatmap v2.18.0. **Spatial distributions and gene expression profiles.** Cell distributions and gene expression were examined along the corticomedullary axis using the distance coordinates stored in the processed Xenium objects. ICA, ICB, and PC distributions were summarized as cell counts in 100-μm intervals. SCT-normalized expression of *Avpr2*, *Aqp2*, *Calb1*, and *Trpv5* was averaged within 250-μm intervals for each sample, followed by equal-weight averaging across biological replicates. Combined expression profiles included intervals containing cells in both samples.

#### KPMP mouse and human whole-kidney snRNA-seq atlases

Whole-kidney snRNA-seq data from normal mouse kidneys (reference and sham-operated) and healthy human reference kidneys were obtained from the KPMP Atlas v2 (https://atlas.kpmp.org/). Nuclei in a reference cell state and annotated as CNT or CNT-PC were retained and integrated across donors with Seurat. Sub-clusters with DCT, collecting duct, IC or other contaminating signatures, predominantly medullary origin, single-donor dominance or low UMI counts were removed. This yielded 2,149 mouse nuclei from 31 mice and 12,559 human nuclei from 59 donors. CNT1, CNT_T_ and CNT2 labels were transferred from CNT nuclei of our distal-enriched dataset to each whole-kidney nucleus by Seurat anchor-based label transfer (canonical correlation analysis, 10 dimensions), using one-to-one orthologs for human. Transfer features were restricted to reference pairwise markers between CNT states that were detected in at least 5% of query nuclei. Prediction scores were averaged over each nucleus’s 30 nearest neighbors, and each nucleus was assigned the state with the highest smoothed score. To identify transitional nuclei in human whole-kidney data independent of clustering, nuclei in the bottom and top 15% of the smoothed CNT2-minus-CNT1 transfer score were defined as CNT1 and CNT2 poles. Pole-specific genes were identified by differential expression in one random half of the pole nuclei and confirmed in the other. Each nucleus was then scored for CNT1-pole and CNT2-pole gene expression, scaled from 0 (opposite pole) to 1 (own pole) and smoothed over 30 nearest neighbors. A two-dimensional Gaussian mixture model was fitted to these scores, with the number of components (1–5) selected by the Bayesian information criterion. Components whose means had both scores between 0.25 and 0.75 were designated transitional (CNT_T_). Nuclei were assigned to a group when the summed posterior probability of its components was at least 0.8 and otherwise classed as uncertain. As a positive control, the procedure was also applied to the distal-enriched reference dataset CNT nuclei without their annotations, using cross-replicate transfer scores. Code development for this section was assisted by Claude Code (Anthropic) under author supervision; all analyses were designed, verified and interpreted by the authors.

## Results

### Immunostaining identifies AQP2-negative and AQP2-positive regions along the connecting tubule

Prior work by other groups suggested that the CNT comprises discrete subsegments in rodents, but these were variably categorized according to pathways for water transport or aldosterone sensitivity.^17,22,23^ To examine this organization, we performed immunostaining in microdissected kidney tubules (Fig. 1A). Consistent with previous reports^17,22^ the CNT appeared to be segmented into AQP2-negative and AQP2-positive regions.

**Figure 1.**
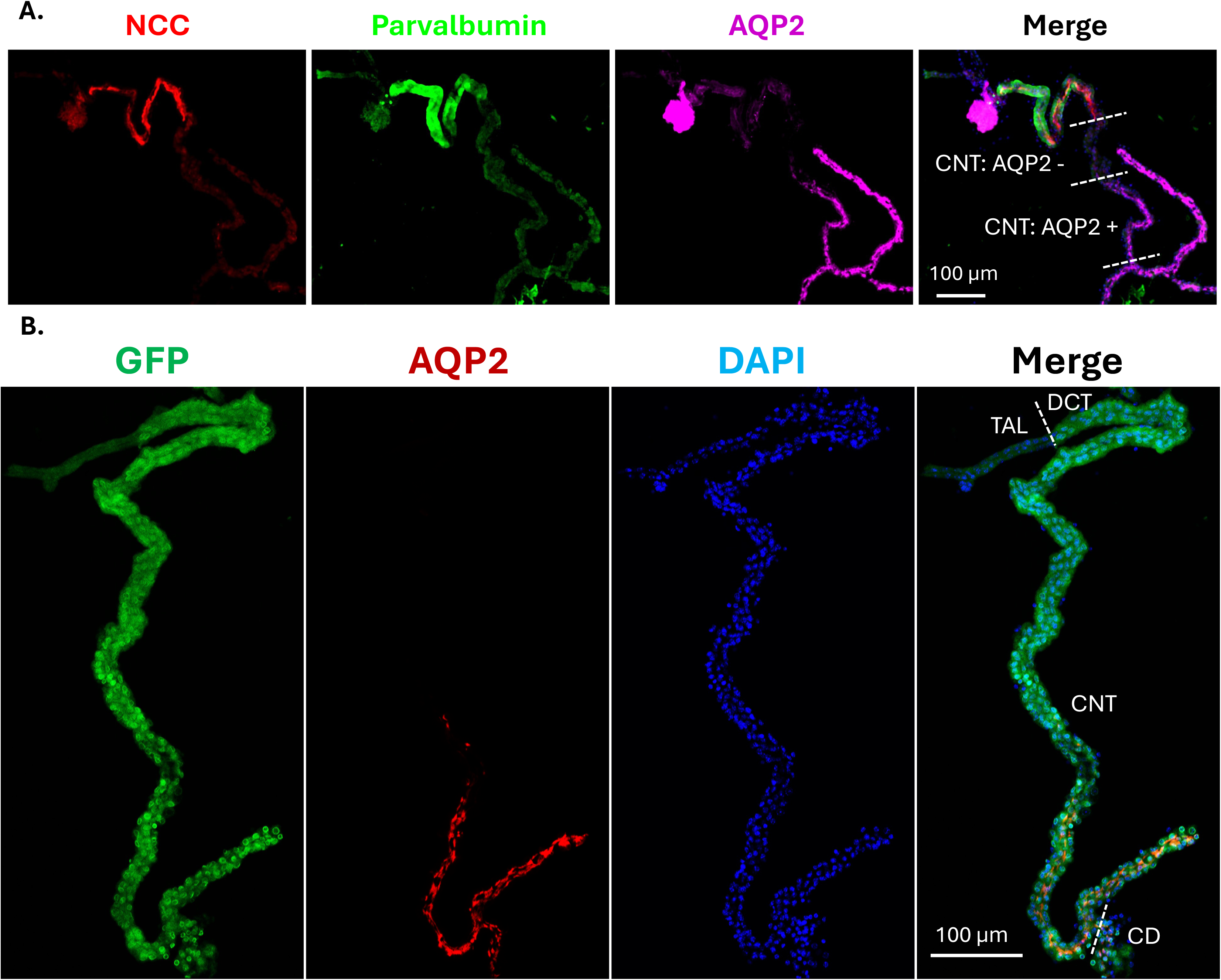
Immunostaining reveals regional AQP2 expression along the CNT and nuclear GFP labeling in Calb1-Cre-INTACT mice. (A) Representative microdissected kidney tubule from wild-type C57BL/6 mouse stained for NCC (red), parvalbumin (green), and AQP2 (magenta). Individual channels and the merged image identify the DCT and adjacent CNT regions with low or undetectable versus prominent AQP2 staining. Dashed white lines delineate the indicated regions. (B) Representative microdissected tubule from a Calb1-Cre-INTACT mouse stained for GFP (green) and AQP2 (red), with DAPI counterstaining of nuclei (blue). GFP labeling of the nuclear membrane is present along the DCT, CNT, and CD profiles indicated in the merged image. Dashed lines mark the annotated TAL-DCT and CNT-CD junctions. The constitutive Cre-dependent reporter labels cells with a history of *Calb1* promoter activity. AQP2, aquaporin-2; CD, collecting duct; CNT, connecting tubule; DCT, distal convoluted tubule; GFP, green fluorescent protein; INTACT, isolation of nuclei tagged in specific cell types; NCC, Na-Cl cotransporter; TAL, thick ascending limb.

Characterization of the CNT is limited by the lack of markers that uniquely distinguish CNT cell populations and by the low abundance of CNT cells, estimated to represent ∼3% of kidney epithelial cells.^9^ To overcome this limitation, we used our established INTACT-based approach to enrich for discrete kidney cell populations.^27,31^ Because *Calb1* has previously been used as a canonical marker of CNT cells in single-cell transcriptomic studies,^9^ we crossed Calb1-Cre mice with the INTACT reporter line, generating mice with Cre-dependent GFP labeling of the nuclear envelope.

Although Calb1 expression in the adult distal tubule is predominantly located in DCT2 and CNT cells^32^, GFP labeling in Calb1-Cre-INTACT mice extended beyond these populations and included labeling along the entire DCT-CNT-CD axis including IC cells (Fig. 1B, Supplemental Fig. 1-2). We attribute this broader labeling to the use of a constitutive Calbindin-Cre line, as previously reported^33^. This strategy provided access to a broad population of distal tubule cells for analysis. GFP-positive nuclei were subsequently enriched by fluorescence-activated nuclei sorting and subjected to snRNA-seq, as previously described.^27^

Calb1-Cre–INTACT mice had normal plasma electrolyte concentrations (Supplemental Table 2).

### Distal tubule-enriched snRNA-seq recapitulates established DCT cell populations and captures all major distal epithelial populations

snRNA-seq of GFP-enriched nuclei from Calb1-Cre-INTACT mice captured multiple epithelial populations of the distal tubule. Unsupervised clustering identified transcriptionally distinct groups, which were visualized using uniform manifold approximation and projection (UMAP, Fig. 2A). DCT populations were identified by expression of *Slc12a3*, encoding the Na-Cl cotransporter NCC (Fig. 2B), and further classified using the transcriptional signatures described in our previous DCT-enriched snRNA-seq study.^27^ The Calb1-Cre-INTACT dataset recapitulated the major features distinguishing DCT1 from DCT2: DCT1 clusters showed prominent expression of *Egf* and *Umod*, whereas DCT2 cells expressed calcium transport–associated genes, including *Calb1*, *Slc8a1*, *S100g*, and *Trpv5* (Supplemental Fig. 3A). The remaining populations were initially annotated using the canonical marker panel described by Chen et al. (Figure 1C, Supplemental Fig. 3B).^9^ Three clusters were identified as CNT populations based on expression of *Calb1* (calbindin-D28K; Fig. 2D) and *Scnn1g* (γ-ENaC; Fig. 2E), together with low or undetectable *Slc12a3* expression. Note that *Calb1* expression in the UMAP (Fig. 2D) reflects expression by mature kidneys in which *Calb1* is largely restricted to DCT2 and CNT cells.^33^ . Although *Aqp2* has been suggested as a gold-standard CNT marker,^9^ its expression varied across CNT clusters, together with that of *Trpv5* and *Hsd11b2* (Fig. 2F-H). Within the UMAP embedding, the cluster nearest DCT2 showed prominent *Trpv5* expression and low or undetectable *Aqp2*, whereas the cluster nearest to the collecting duct principal cells (PC-CD) displayed the reciprocal pattern (Aqp2^+^/TRPV5^-^). These populations were designated CNT1 and CNT2, respectively, consistent with a previously proposed nomenclature.^22^A third cluster occupied an intermediate position and displayed features of both populations. We designated this population as transitional CNT (CNT_T_) to indicate its intermediate phenotype and suggest that it may reflect a regulated cell state that is transitional between CNT1 and CNT2.

**Figure 2.**
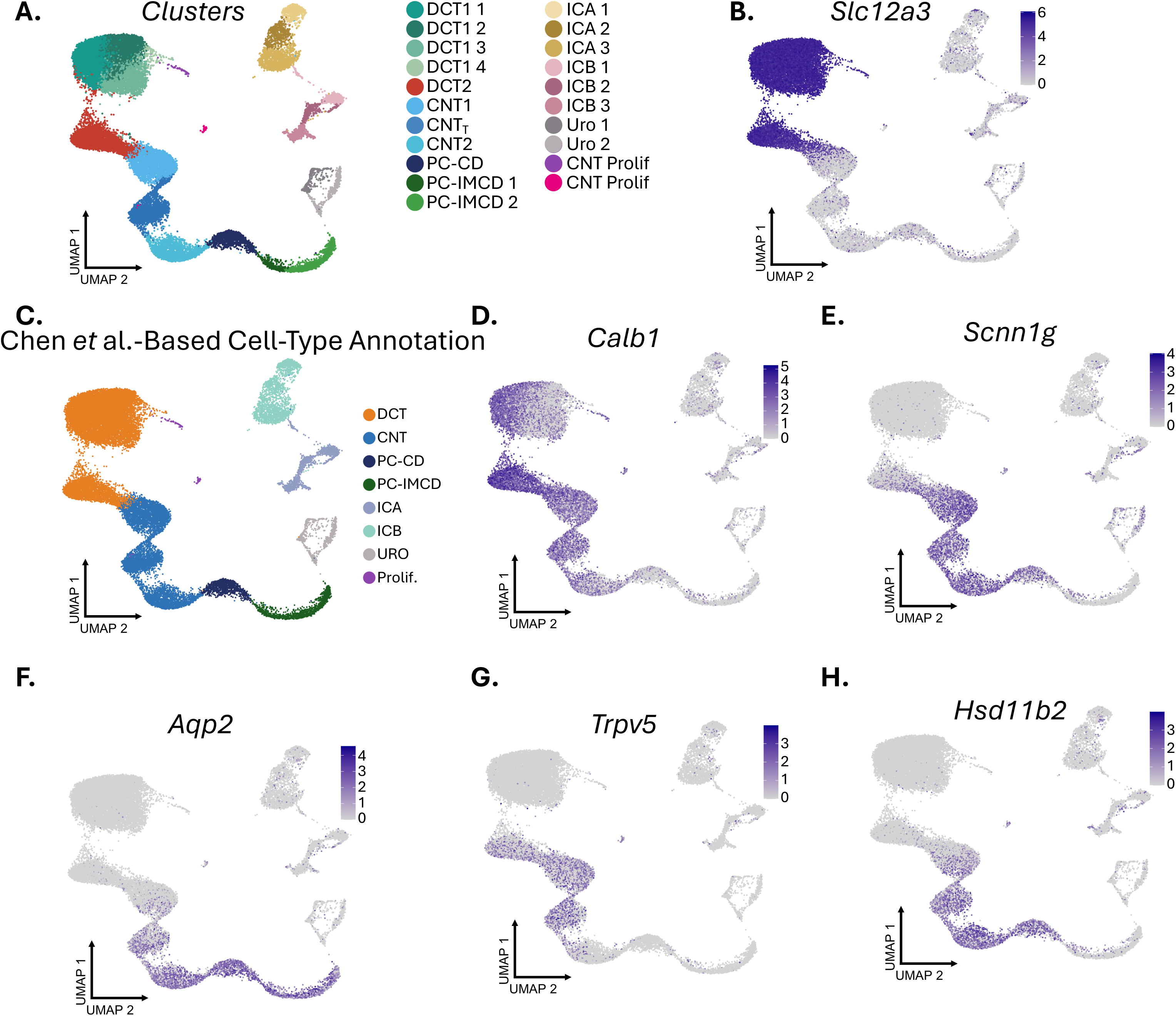
Distal tubule enrichment resolves transcriptionally distinct epithelial populations. Single-nucleus RNA sequencing was performed on GFP-enriched nuclei isolated from the kidneys of two male Calb1-Cre-INTACT mice. (A) Uniform manifold approximation and projection (UMAP) of the combined dataset, colored by annotated cluster. Each point represents one nucleus. DCT, CNT, CD, IC, urothelial (URO), and proliferating (prolif) populations are indicated. (B) Expression of *Slc12a3* (NCC). (C) The same nuclei grouped into major epithelial populations using the canonical marker panel described by Chen et al. ^9^. (D-G) Expression of *Calb1* (calbindin-D28k; D), *Scnn1g* (γ-ENaC; E), *Aqp2* (aquaporin-2; E), *Trpv5* (F), and *Hsd11b2* (11β-hydroxysteroid dehydrogenase type 2; G) projected onto the same UMAP. Color intensity represents normalized transcript expression, with gene-specific scales. *Calb1* transcript abundance reflects expression at the time of sampling in adult mice. (H). CNT1 and CNT2, connecting tubule populations 1 and 2; CNT_T_, transitional CNT; DCT1 and DCT2, distal convoluted tubule 1 and 2; ENaC, epithelial sodium channel; ICA and ICB, type A and type B intercalated cells; PC-CD, collecting duct principal cells; PC-IMCD, inner medullary collecting duct principal cells.

Principal cells of the collecting duct (PC-CD), including those from the cortical collecting duct (CCD) and outer medullary collecting duct (OMCD), were defined by expression of *Aqp2* (Fig. 2F) and *Scnn1g* (Fig. 2E) but low or undetectable *Calb1* expression (Fig. 2D). PC cells of the inner medullary collecting duct (IMCD) were characterized by expression of *Aqp2* and *Slc14a2* (UT-A1) (Supplemental Fig. 3A). IC were identified by expression of *Slc4a1* (anion exchanger 1) or *Slc2ca4* (pendrin) (Supplemental Fig. 3A-B). Two additional minor populations were identified: urothelial cells (URO) expressing *Upk1b* and proliferating DCT and CNT cells expressing the cell-cycle marker *Top2a* (Supplemental Fig. 3A), consistent with previous reports.^29,30^

Although all major distal tubule cell types were identified, owing to the broad developmental Calb1 expression, subsequent analyses focused specifically on cell populations that express Calb1 during adulthood; the remaining annotated distal tubule populations were grouped as “others.” The full set of annotated cell-type clusters is provided in the Supplemental Material.

### Distinct CNT populations exhibit graded expression of calcium, sodium, and water transport genes

Differential gene (DEG) expression analysis revealed distinct expression profiles among three CNT clusters sharing *Calb1* and *Scnn1g* expression together with low or undetectable *Slc12a3* (Figure 3A). Top DEGs are shown in supplemental figure 4.

**Figure 3.**
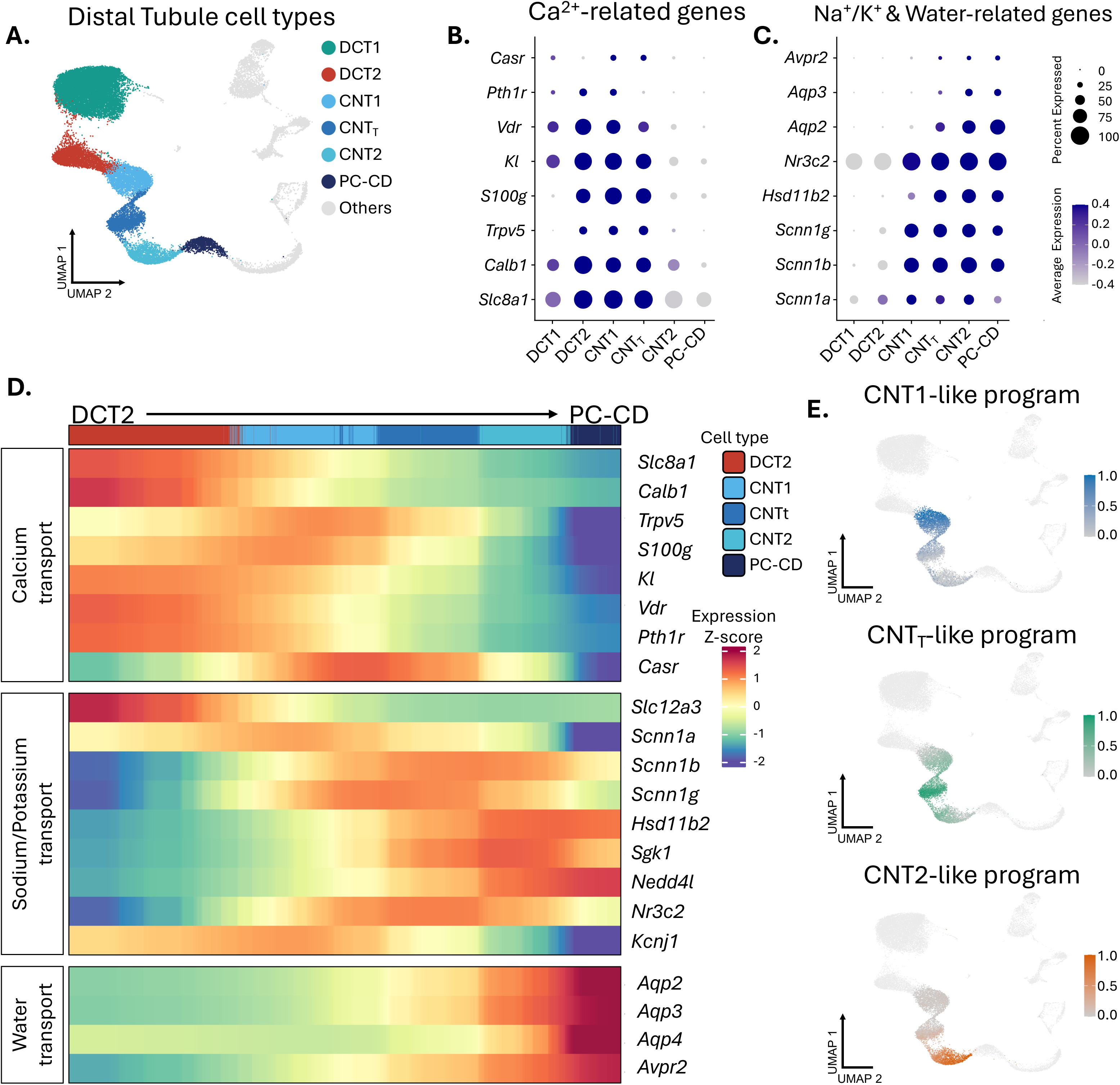
Calcium, sodium, and water transport-associated expression varies across CNT populations. (A) UMAP highlighting DCT1, DCT2, CNT1, CNTT, CNT2, and PC-CD populations in the Calb1-Cre-INTACT single-nucleus RNA sequencing dataset. Other annotated populations are shown in gray. (B and C) Dot plots of genes associated with calcium transport (B) and sodium/potassium transport, mineralocorticoid signaling, and water transport (C). Dot size indicates the percentage of nuclei with detectable expression, and color indicates gene-wise scaled average expression within each population. (D) Pseudotime-ordered heatmaps of selected transport-associated genes in DCT2, CNT1, CNT_T_, CNT2, and PC-CD nuclei, with the trajectory rooted in DCT2. Log-normalized expression was smoothed across ordered nuclei and standardized to gene-wise z-scores. The upper annotation bar identifies the cell types. Calcium-associated expression is prominent toward DCT2 and CNT1, whereas water-associated expression increases toward CNT2 and PC-CD, with sodium transport– associated genes expressed across the CNT. (E) Per-nucleus usage of the CNT1-like, CNT_T_-like and CNT2-like programs, shown on the UMAP in (A). The programs came from consensus non-negative matrix factorization (cNMF; K = 5) of 6,892 CNT nuclei from vehicle-treated mice, using all genes. Usage is normalized to sum to 1 across the five programs in each nucleus. Non-CNT cells are shown in grey. CNT, connecting tubule; CNT_T_, transitional CNT; DCT, distal convoluted tubule; PC-CD, collecting duct principal cells; UMAP, uniform manifold approximation and projection.

CNT1 was distinguished by prominent expression of genes associated with calcium transport, including *Slc8a1* (NCX1), *Calb1* (calbindin-D28K), *Trpv5*, *S100g* (calbindin-D9K), *Kl* (Klotho), *Vdr* (vitamin D receptor), and *Pth1r* (parathyroid hormone receptor 1) (Fig. 3B). CNT1 and DCT2 share a core calcium transport gene signature but exhibit distinct transcriptional profiles, suggesting distinct functional specialization. CNT_T_ retained substantial expression of several calcium transport-associated genes, whereas their expression was markedly lower in CNT2.

Genes associated with sodium and water transport, as well as aldosterone-sensitivity, were more broadly distributed. *Scnn1b* and *Scnn1g*, encoding the β- and γ-subunits of ENaC, and *Nr3c2*, encoding the mineralocorticoid receptor, were expressed across all three CNT populations (Fig. 3C). In contrast, *Hsd11b2*, encoding the enzyme that limits glucocorticoid activation of the mineralocorticoid receptor, showed low expression in CNT1 and more prominent expression in CNT_T_ and CNT2. Water transport genes also distinguished these populations: *Aqp2* expression became apparent in CNT_T_ and was higher in CNT2 and PC-CD, while *Aqp3* was predominantly expressed in CNT2 and highest in PC-CD. CNT_T_ therefore combined retention of calcium transport–associated genes with increased expression of *Hsd11b2* and *Aqp2*, supporting its designation as an intermediate transcriptional phenotype.

To examine these patterns across the DCT2-CNT-PC-CD populations, we constructed a pseudo-spatial arrangement, starting from DCT2 and visualized selected genes in ordered heatmaps (Fig. 3D). The resulting profiles showed overlapping expression patterns rather than uniformly abrupt boundaries. *Calb1* and *Slc8a1* were most prominent toward the DCT2 end of the trajectory, whereas *Trpv5* and *S100g (*calbindin-D_9k_) reached their strongest expression within CNT1.

Regarding sodium transport machinery, *Scnn1a*, encoding α-ENaC, was also most prominent in CNT1 and declined modestly across CNT_T_ and CNT2. By comparison, *Scnn1b*, *Scnn1g*, and *Hsd11b2* showed greater expression toward the CNT_T_-CNT2 portion of the trajectory. Increasing expression of *Aqp2*, *Aqp3*, *Aqp4*, and *Avpr2* also characterized the progression toward PC-CD. This transcriptional ordering highlighted a gradual shift from calcium transport–associated expression toward water transport–associated expression. Additionally, expression of Hsd11b2, responsible for conferring aldosterone sensitivity, largely aligned with that of water-transport genes. Module scores for calcium, sodium, and water transport gene cassettes further supported these expression patterns when visualized on the UMAP (Supplemental Fig. 5).

To determine whether the CNT clusters reflect underlying transcriptional programs independent of clustering, we applied consensus non-negative matrix factorization (cNMF) to CNT nuclei without reference to cluster labels.^28^ Three of the six identified programs corresponded to the annotated clusters (Fig. 3E). A CNT1-like program predominated in CNT1, a CNT_T_-like program in CNT_T_ , and a CNT2-like program in CNT2. Program usage was graded rather than discrete, with CNT1-like and CNT2-like usage extending into CNT_T_, consistent with the gradual shift in gene expression described above.

To examine whether transcriptional heterogeneity was reflected at the protein level, we performed immunofluorescence microscopy on perfusion-fixed kidneys. CNT and collecting duct profiles were distinguished using anatomical characteristics. Within anatomically identified CNT profiles, regions with strong calbindin-D28k (Calb1) staining and low or undetectable AQP2 were consistent with CNT1, whereas regions with low but detectable calbindin-D28k and robust AQP2 were assigned to CNT2 (Fig. 4A). Additional profiles illustrate similar regional differences in the relative abundance of both proteins (Fig. 4B-D).

**Figure 4.**
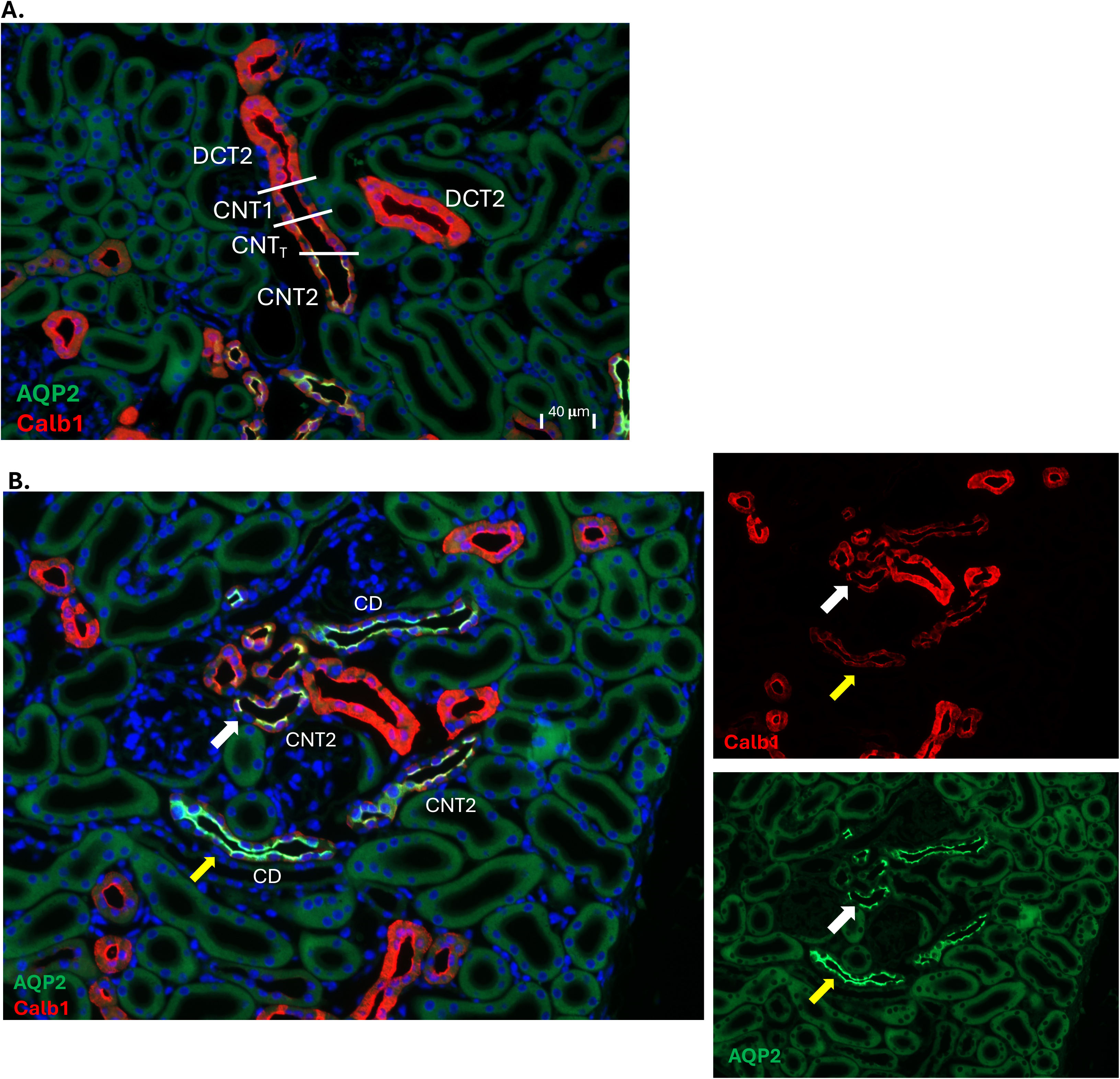
Calbindin-D28K and AQP2 exhibit heterogeneous expression along cortical distal tubules. Representative immunofluorescence images of perfusion-fixed mouse kidney sections stained for calbindin-D28k (red) and AQP2 (green), with DAPI nuclear counterstaining (blue). (A) A longitudinal tubular profile illustrates changes in the relative abundance of calbindin-D28K and AQP2 across regions annotated as DCT2, CNT1, CNT_T_, and CNT2. White lines mark the indicated regions. Calbindin-D28K staining is prominent in the DCT2/CNT1 regions, whereas AQP2 becomes more prominent toward the CNT2 region. (B) Additional cortical profiles illustrating variation in calbindin-D28K and AQP2 staining in regions labeled CNT2 and CD. White and yellow arrows identify corresponding tubular profiles across the images. Segment labels indicate the proposed correspondence of the anatomical and staining features with the transcriptionally defined populations. AQP2, aquaporin-2; CD, collecting duct; CNT, connecting tubule; CNT_T_, transitional CNT; DCT2, distal convoluted tubule 2.

This work suggested that pathways for calcium reabsorption were largely expressed proximal to pathways for water transport, but the segmental distinctions were not abrupt. TRPV5 and AQP2 staining provided complementary evidence for regional differences in transport protein expression. These proteins are predominantly localized to different tubular regions, with little apparent overlap in the profiles examined (Fig. 5A-C). Together with the transcriptomic findings, these observations support regional separation of the molecular machinery associated with calcium entry and water transport along the CNT.

**Figure 5.**
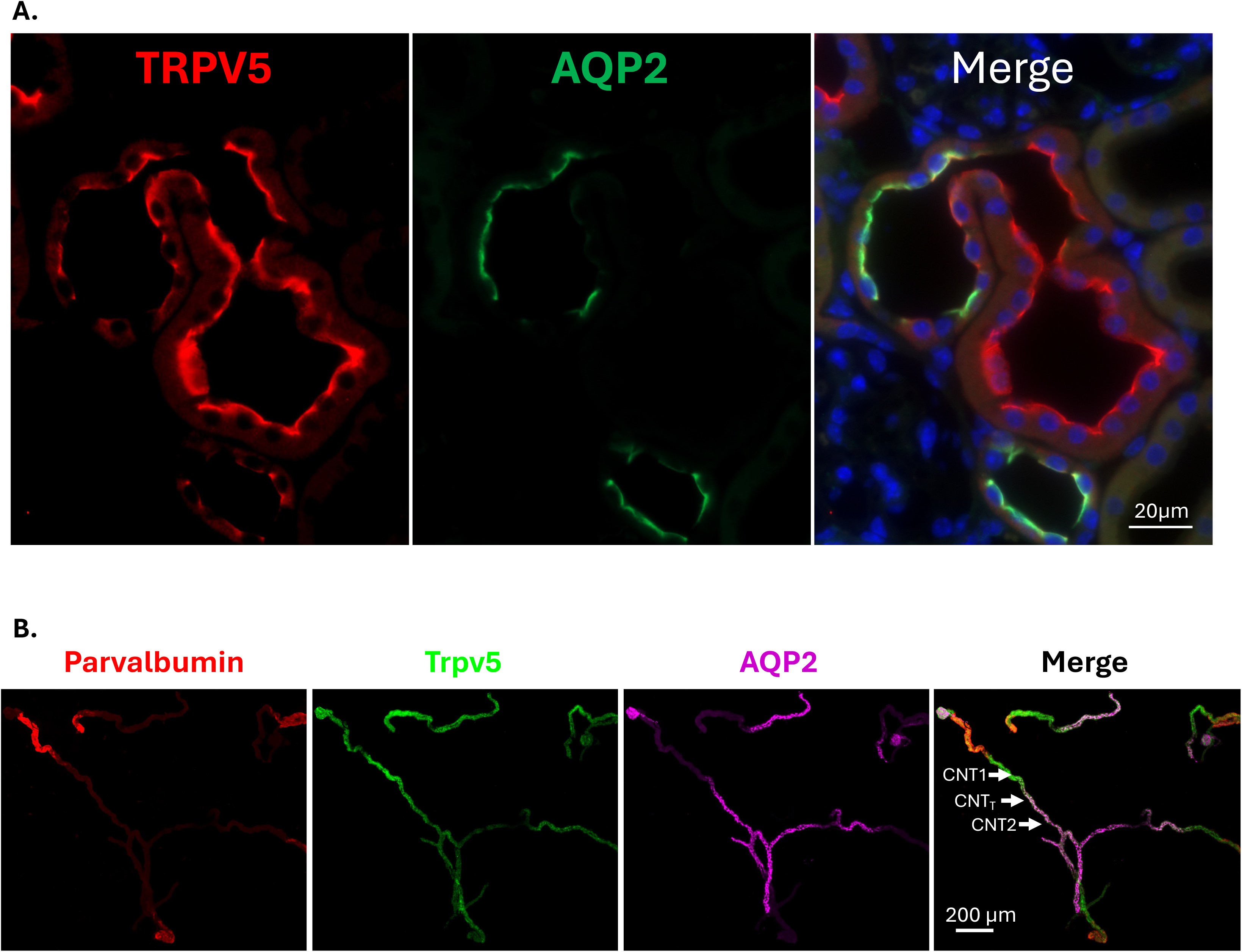
TRPV5 and AQP2 display regional differences in distal tubule protein expression. (A) Representative mouse kidney section stained for TRPV5 (red) and AQP2 (green). Nuclei are counterstained with DAPI (blue). (B) Representative microdissected kidney tubule stained for parvalbumin (red), TRPV5 (green), and AQP2 (magenta). The merged images illustrate the change from TRPV5-predominant to AQP2-predominant staining along the CNT. Arrows identify regions annotated as CNT1, CNT_T_, and CNT2. AQP2, aquaporin-2; CNT, connecting tubule; CNT_T_, transitional CNT; TRPV5, transient receptor potential cation channel subfamily V member 5.

Expression of these and other relevant genes across all major distal tubule populations is shown in Supplemental Figs. 6 and 7.

### CNT populations exhibit distinct nephron- and ureteric-associated transcriptional signatures

To explore whether CNT heterogeneity reflected developmental origin, we examined nephron- and ureteric-lineage-associated markers reported by Ransick et al.^34^ The nephron-associated marker *Slc2aS* was preferentially expressed in DCT2, CNT1, and CNT_T_, whereas CNT2 shared ureteric-associated expression features with PC-CD, including detectable *Ache* expression (Supplemental Fig. 8). Comparison with the broader segment-specific marker panel is shown in Supplemental Fig. 9). These patterns suggest that CNT heterogeneity may reflect differences in developmental origin, although lineage tracing would be required to establish the ancestry of these populations and locate the nephron– ureteric junction.

### Spatial transcriptomics supports an ordered continuum of CNT populations

To determine whether CNT transcriptional heterogeneity corresponded to a spatially ordered continuum or an interspersed cellular arrangement, we analyzed a publicly available Xenium dataset from two male mouse kidneys.^29^ DCT, CNT, collecting duct principal cells, intercalated cells, and urothelial cells were selected to approximate the epithelial populations represented in the Calb1-Cre–INTACT dataset. These populations were re-clustered and annotated using the described markers available in the Xenium panel to identify major distal tubule cell types. All remaining kidney cell types were grouped as “Others.” CNT populations were further classified using expression patterns from the Calb1-Cre–INTACT dataset, with particular attention to *Trpv5* and *Aqp2*.

Despite the limited gene coverage of the Xenium panel, pairwise expression differences among CNT1, CNT_T_, and CNT2 correlated with those in the snRNA-seq reference in both replicates (Spearman ρ = 0.58-0.83; 74-86% concordant in direction), although their magnitude was attenuated (Supplemental Fig. 10A-B). Likewise, reference-derived cNMF programs projected onto Xenium cells supported the ordering CNT1 < CNT_T_ < CNT2 in both samples (Supplemental Fig. 11). The CNT_T_-like program was less distinct in Xenium than in the reference, which we discuss further below.

Spatial mapping and regional quantification showed that DCT1, DCT2, and all three CNT populations were almost exclusively cortical (Fig. 6A-C; Supplemental Figs. 12 and 13). Enlarged views of the same cortical region, shown with and without intercalated cells, suggested a DCT2–CNT1–CNT_T_–CNT2–PC-CD arrangement (Fig. 6B and Supplemental Fig. 13C).

**Figure 6.**
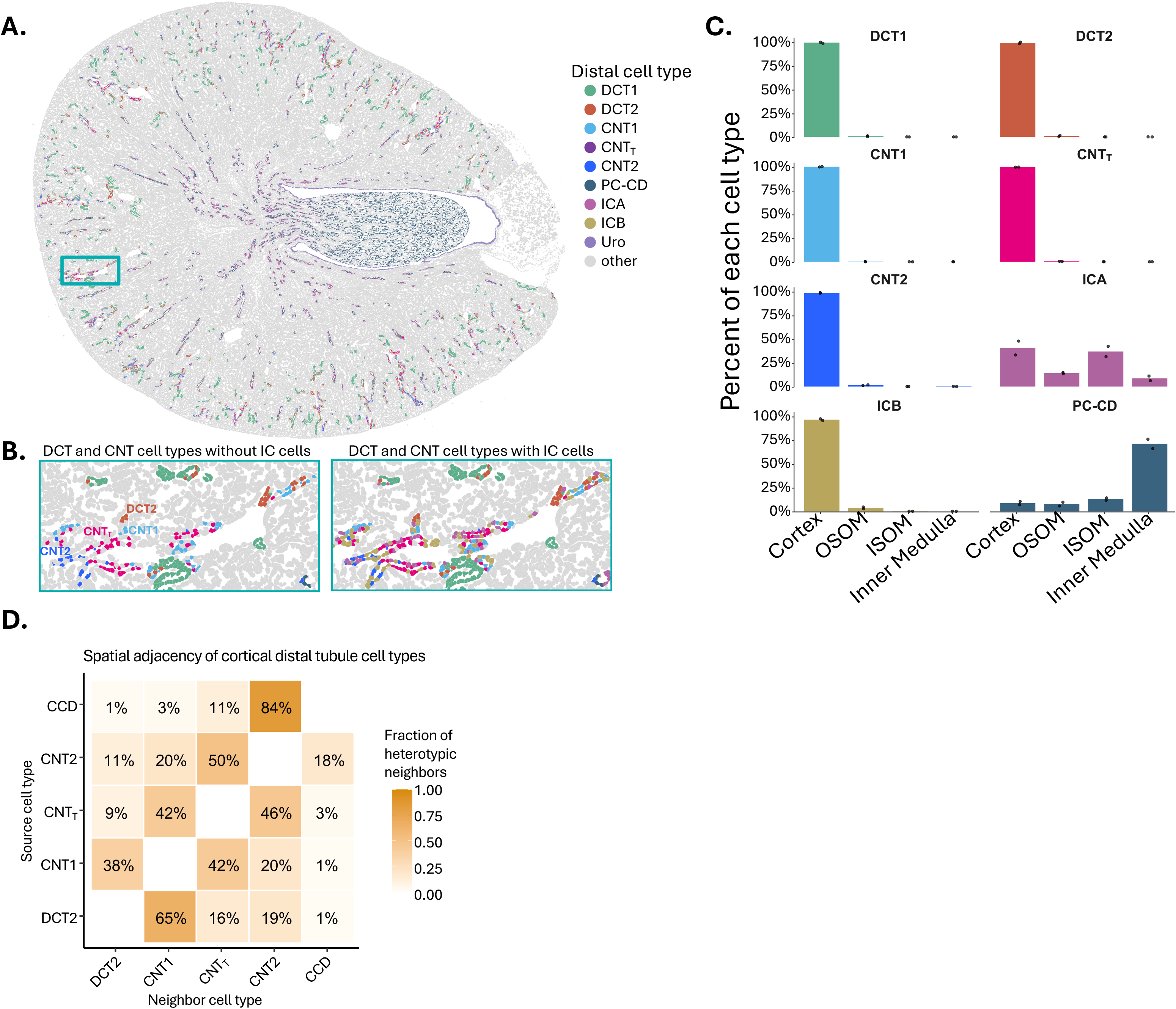
Spatial transcriptomics supports an ordered arrangement of CNT populations in the mouse kidney. A publicly available Xenium dataset from two male mouse kidneys was reanalyzed ^29^. Distal tubule epithelial populations were re-clustered and annotated using markers available in the Xenium panel and expression patterns from the Calb1-Cre–INTACT dataset. (A) Spatial map of sample 1, with annotated populations colored as indicated and other kidney cell types shown in gray. The outlined cortical region is enlarged in B. (B) The same region displayed without (left) and with (right) intercalated cells highlighted, showing the arrangement of DCT and CNT populations within tubular profiles. (C) Regional distribution of each annotated population across the cortex, outer stripe of the outer medulla (OSOM), inner stripe of the outer medulla (ISOM), and inner medulla. Percentages were calculated within each cell type and sample; bars show the mean of two samples, and points represent individual samples. (D) Heterotypic spatial neighborhood analysis restricted to cortical DCT2, CNT1, CNT_T_, CNT2, and PC-CD. Neighbors were defined by a cell-centroid distance of 20 μm or less. For each source population (rows), colors and values indicate the fraction and percentage of heterotypic neighbor links assigned to each population (columns). Homotypic links were excluded and are shown as blank diagonal entries, IC were also excluded. Results combine both samples. Panels C-D include two biological replicates. CNT, connecting tubule; CNT_T_, transitional CNT; DCT, distal convoluted tubule; ICA and ICB, type A and type B intercalated cells; PC-CD, collecting duct principal cells; CCD, cortical collecting duct, URO, urothelial cells.

To assess this organization quantitatively, we analyzed heterotypic neighbors, defined as cells of different annotated types within a 20-μm radius. Because intercalated cells predominated among CNT heterotypic neighbors in the broader analysis, we examined relationships among cortical DCT2, CNT1, CNT_T_, CNT2, and PC-CD cells. DCT2 most frequently neighbored CNT1, CNT_T_ neighbored both CNT1 and CNT2, and principal cortical CD cells predominantly neighbored CNT2, supporting the proposed spatial ordering (Fig. 6D). Individual-sample results and the broader analysis of all distal populations are shown in Supplemental Fig. 14. Sample 2 contained less cortical tissue, limiting its anatomical coverage.

Pseudo spatial-ordered heatmaps incorporating both samples and rooted in DCT2 recapitulated the expression patterns observed by enriched snRNA-seq (Supplemental 15). Additionally, we mapped gene expression along the corticomedullary axis, with zero marking the outer-inner medullary boundary. Across both samples, *Calb1* and *Trpv5* expression declined from cortical toward medullary positions, whereas *Aqp2* and *Avpr2* increased (Supplemental Fig. 16A). Cell counts along the same axis placed principal cells predominantly in the inner medulla and type B intercalated cells in the cortex. Type A intercalated cells extended across cortical and medullary regions (Supplemental Fig. 16B).

### The CNT gradient with a transitional population is recapitulated in humans

To determine whether the CNT1-to-CNT2 organization is conserved in human, we analyzed healthy reference nuclei from the KPMP Atlas v2.^35^ We retained 12,559 high-confidence CNT nuclei, comprising cells annotated as CNT or CNT-PC in the atlas (Fig. 7A). We then transferred CNT labels to each human nucleus by Seurat anchor-based label transfer, using our distal-enriched mouse snRNA-seq dataset as the reference, mapped to human genes through one-to-one orthologs. Transferred labels placed human CNT nuclei along a continuous gradient, from nuclei resembling mouse CNT1 (hereafter CNT1) to nuclei resembling mouse CNT2 (hereafter CNT2) at the CNT-PC pole (Fig. 7B).

**Figure 7.**
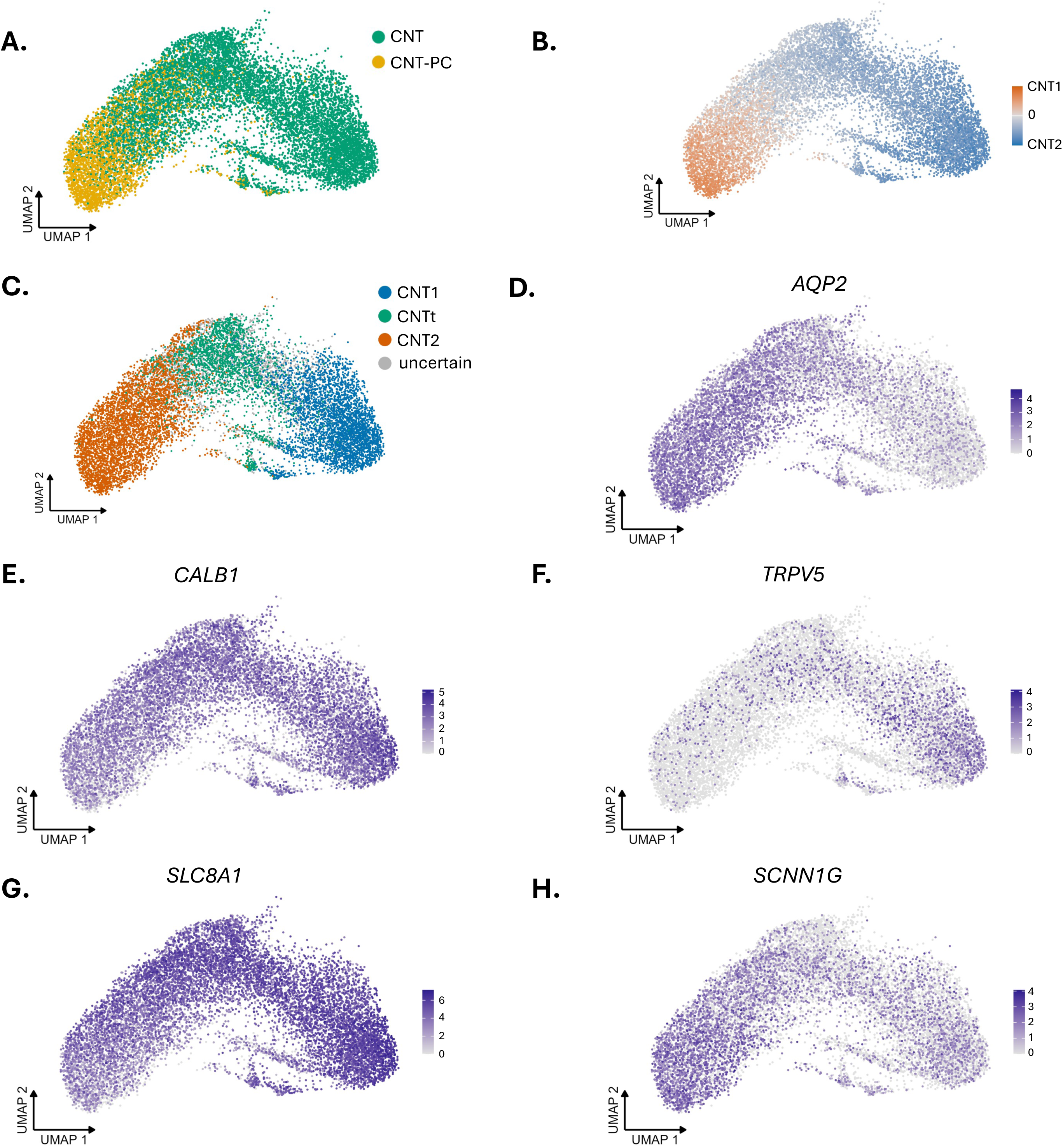
Human CNT profiles exhibit a continuum of calcium-and water transport–associated expression. Human kidney transcriptomic data from the Kidney Precision Medicine Project (KPMP) Atlas v2,^52^ comprising 12,559 healthy-reference CNT nuclei from 59 donors. (A) Uniform manifold approximation and projection (UMAP) of human CNT showing original KPMP subclass labels: CNT and CNT-PC. (B) CNT1-to-CNT2 transfer score from mouse CNT labels (CNT1-CNT_T_-CNT2) transferred from our distal-enriched snRNA-seq reference. The score is the CNT2 prediction score minus the CNT1 prediction score, averaged over 30 nearest neighbors. Positive (orange) values are CNT2-like and negative (blue) values are CNT1-like. (C) Unsupervised classification of human nuclei into CNT1 (29.0%), CNT_T_ (16.5%) and CNT2 (35.4%). Classes came from a Gaussian mixture model fitted to two pole-derived expression scores. Nuclei with a grouped posterior probability below 0.8 are uncertain (19.1%, grey). (D–H) Log-normalized expression of AQP2 (D), CALB1 (E), TRPV5 (F), SLC8A1 (G) and SCNN1G (H). Purple intensity indicates transcript expression, with gene-specific scales. The expression maps illustrate variation in calcium-and water-associated genes across the CNT continuum, together with expression of the sodium channel subunit SCNN1G. CNT, connecting tubule; CNT-PC, connecting tubule-principal cell population; CNT_T_, transitional CNT.

Because mouse CNT_T_ cells co-express CNT1 and CNT2 features, we searched for human nuclei with this property without relying on clustering. Human CNT1- and CNT2-associated gene programs were derived from nuclei at the two poles of the gradient, and each nucleus was scored for both. Mixture modelling of these scores resolved a population with intermediate expression of both programs (16.5% of nuclei; 15.2–18.1% across alternative pole definitions), which occupied a contiguous region between the CNT1 and CNT2 poles (Fig. 7C; Supplemental Fig. 17A). Applied to mouse data without annotations, the same approach recovered 96% of CNT_T_ cells with 92% precision (Supplemental Fig. 17B). In contrast, direct transfer of the mouse CNT_T_ label did not identify a corresponding human population (2–8% of nuclei, without a defined location) (Supplemental Figs. 18A-C), and human nuclei showed no preferential use of the mouse CNT_T_-specific program (Supplemental Figs. 18D-F). Human CNT therefore contains a transitional population defined by co-expression of CNT1 and CNT2 programs, but not the CNT_T_-specific signature observed in mouse. Canonical markers followed the expected distribution: *TRPV5* were enriched toward the CNT1 pole, *AǪP2* and *SCNN1G* toward the CNT2 pole, and *CALB1* and *SLC8A1* throughout (Fig. 7D–H).

Lastly, the intermediate character of CNT_T_ was consistent across datasets, including a whole-kidney mouse snRNA-seq atlas from KPMP v2 (Supplemental Figs. 18G-I) In this whole-kidney dataset, as in human, CNT nuclei did not resolve into a separate CNT_T_ cluster, which emerged only in our distal-enriched dataset. CNT_T_ was therefore assigned by label transfer from the distal-enriched reference (Supplemental Fig. 18). Across mouse distal-enriched snRNA-seq, two Xenium replicates, mouse whole-kidney snRNA-seq and human, most CNT1-versus-CNT2 differentially expressed genes were expressed in CNT_T_ at levels between their corresponding CNT1 and CNT2 (68–96%) (Supplemental Fig. 19A). At the single-cell level, CNT_T_ cells formed a single population between CNT1 and CNT2 cells, most of which co-expressed CNT1 and CNT2 genes (Supplemental Fig. 19B). Xenium replicate 2 was not interpretable at this level because CNT1 and CNT2 cells were insufficiently separated.

## Discussion

Our integrated findings support a model in which the rodent and human CNT forms part of a gradual transcriptional continuum linking the DCT to the collecting duct. Along this continuum, gene expression shifts from calcium transport-associated pathways toward water transport-associated pathways, with 11-β-hydroxysteroid dehydrogenase, which confers aldosterone sensitivity, arising in same regions as regulated water flux. These findings help reconcile longstanding uncertainty about CNT identity and its boundaries. Balzer et al highlighted this ambiguity, noting that populations designated as CNT in their datasets closely resembled cells classified as DCT2 elsewhere.^36^ Yang and colleague also noted the similarity between what has been classified as DCT2 and portions of the CNT.^17^ Confusion also reflects the fact that many studies focus on selected aspects of solute transport, water permeability, or hormonal regulation; this means that different and sometimes conflicting CNT classifications exist.^17,19–23,37^ By applying enrichment strategy to studies of the DCT and TAL, we clarified the nature of specialized epithelial populations.^27,31^ Here, nuclear enrichment was central to characterizing less abundant CNT cells, enabling us to resolve expression differences that can be obscured in whole-kidney datasets and to place them within a gradual distal tubule continuum.

Identifying and defining biologically distinct cell types is fundamentally a heuristic process that requires evidence beyond transcriptional clustering alone.^38,39^ We therefore complemented our enriched transcriptional analysis with immunostaining of kidney sections, analysis of dissected tubules, independent mouse whole-kidney and human biopsy datasets, and spatial transcriptomics. The concordance of results across techniques supports the presence of distinct CNT cell types and the gradual transition between transport-associated programs; it suggests that an intermediate population is transitional and subject to physiological control. Each approach contributes different strengths and limitations, considered below; together, they provide a framework for interpreting the physiological significance of calcium, sodium, potassium, and water transport organization along the CNT.

Early studies of CNT function used dissected tubules from rabbits and found that the CNT actively absorbs calcium and has very low water permeability, unaffected by arginine vasopressin.^2^ Almeida and Burg later documented electrogenic sodium transport by this segment and noted that transport rates are much higher in the CNT than in the CD under similar conditions.^40^ Yet, while the transitions between DCT, CNT, and CCD are clear and abrupt in rabbits, they are gradual in rodents and humans. Some have suggested that the organization of solute and water transport proteins in rodents and humans is different from that in rabbit.^9^ Chen and colleagues noted these differences in a paper proposing a “Kidney Tubule Epithelial Cell Nomenclature for Single-Cell RNA-Sequencing Studies”,^9^ building on the Standard Nomenclature for Kidney Structures, published by the *Renal Commission of the International Union of Physiologic Sciences*.^41^ Chen and colleagues noted that the expression pattern of transport proteins in rabbits aligns well with morphological divisions. In contrast, areas of NCC, ENaC, TRPM6, TRPV5, and AQP2 expression were suggested to overlap in rodents and humans.^9^

Our distal tubule enriched transcriptomic approach, coupled with dissected tubule and immunocytochemical analysis, allowed us to examine many pathways and regulatory factors simultaneously. Prior work has indicated that distal calcium absorption in rodents is not restricted to the CNT but also takes place along the DCT2.^42,43^ Here, we confirm that calcium reabsorbing genes are expressed along both DCT2 and CNT, although their distribution is not uniform. The sodium-calcium exchanger (*Slc8a1*) and calbindin-D28k are most highly expressed along DCT2, where receptors for vitamin D and parathyroid hormone are also abundant. Surprisingly, the apical calcium channel, *Trpv5*, is most highly expressed more distally, along CNT1 and CNT_T_, locations where the calcium-sensing receptor, and calbindin-D9k are also most highly expressed. The segmental expression pattern suggests that rates of calcium reabsorption peak in the DCT2, where delivered load it highest, but that Trpv5 expression is higher distally, where calcium is absorbed against a steeper electrochemical gradient; in fact, lumen calcium concentrations can decline close to zero along the CNT during extracellular fluid volume depletion.^44^

Notably, there is very little expression of aquaporin-2, -3, or -4 or type 2 vasopressin receptors in the segments that reabsorb calcium. This suggests that calcium is mostly reabsorbed before water is absorbed, an arrangement that should help prevent luminal calcium precipitation. These results, however, suggest that the overall organization of solute and water transport pathways along the CNT of rodents and humans is more like rabbit than often appreciated. Traditional models suggest that the CNT of rabbits lacks aquaporin-2 and therefore exhibits low water permeability.^2^ In rodents, however, the CNT is has been suggested to be water-permeable,^21^ although some have suggested that it comprises distinct water-permeable and water impermeable portions.^19,20,22,37^ The present results confirm that some CNT segments have abundant aquaporin-2 expression, whereas others have low or absent Aqp2 expression (and vasopressin type 2 receptor expression). Thus, calcium and water transport regions appear to be separated spatially in all three species. We suggest, therefore, that *Aqp2* should no longer be considered a ‘gold standard marker’ for rodent CNT cells,^9^ because this gene is expressed by only some CNT cells.

The results regarding sodium transport and related regulatory genes are also illuminating. As expected, ENaC is expressed from the DCT2 through the CNT and into the CD, but the abundance of subunits does not align. While α-ENaC is most highly expressed in the DCT2 and CNT1, sites that are mineralocorticoid-dependent but aldosterone-independent ^45–47^, β-ENaC and γ-ENaC are more highly expressed distally. These are the regions that express abundant *Hsd11b2*, which renders them responsive to aldosterone. As MR stimulation increases α-ENaC but not β-ENaC or γ-ENaC,^48^ the aldosterone induction of α-ENaC along these more distal segments should increase electrogenic sodium absorption. Additionally, the concordance with sites of vasopressin responsiveness should further activate ENaC during volume stress,^49^ thereby maintaining potassium secretion in the face of higher tubule fluid potassium concentration.^50^

CNT_T_ emerged as a discrete cluster only in our distal tubule–enriched mouse snRNA-seq dataset. In mouse and human whole-kidney datasets and in Xenium, CNT cells instead formed a continuous CNT1-to-CNT2 gradient, although cells at the corresponding position retained an intermediate profile. We attribute this difference mainly to resolution. Enrichment focuses the analysis on closely related distal populations, so differences among them become a major source of transcriptional variance. In whole-kidney preparations, variance is dominated by differences between major cell types, CNT cells are sparsely represented, and integration across studies and chemistries can attenuate subtle within-segment variation. In Xenium, the targeted panel lacked most genes that distinguish CNT_T_, so these cells could be placed along the gradient but not resolved by their own signature. In human samples, donor heterogeneity, limited biopsy sampling, and possible species differences may contribute further. Whether a population separates as its own cluster therefore depends on sampling depth, cell number, and reference context, and is not by itself a test of biological identity. For this reason, we do not propose CNT_T_ as a distinct cell type. We use the term to denote cells at an intermediate position that share features of CNT1 and CNT2, along a transition that is gradual rather than abrupt, as it is at the TAL–DCT1 boundary.^51^ This gradual organization may itself be physiologically meaningful. We speculate that transitional cells may adopt more CNT1-like or more CNT2-like programs as physiological demands change. That would shift the effective boundary between the calcium-reabsorbing, water-impermeable portion of the CNT and its aldosterone-sensitive, water-permeable portion. This would be consistent with the vasopressin-induced lengthening of the Aqp2-expressing segment.^19^ It could be tested by lineage tracing, or by measuring the size and composition of the transitional population after perturbing calcium, sodium, potassium, or water balance. Therefore, these findings suggest that transitional states along the distal tubule may be under-resolved in whole-kidney atlases and targeted enrichment can complement atlas-scale efforts.

Based on these results, and the work of others, we propose that the distal tubule model proposed by Chen and colleagues, which includes sodium, calcium and water transport pathways,^9^ be modified (Figure 8). While the sites of transport gene expression in rabbits, shown in their Figure 2 appear secure, calcium transport in rodents seems also to take place across CNT segments with low Aqp2 expression. Additionally, sites of magnesium transport (via Trpm6) and calcium transport are sequentially oriented, much like in the rabbit. Finally, we introduce sites of Hsd11b2 expression, as aligning with Aqp2. In the Kidney Precision Medicine Project database, two types of CNT cell are notated, CNT and CNT-PC, the latter expressing more Aqp2. These cells are quite similar to principal cells of the cortical CD, but in our analysis, cluster separately. The definitions and even the existence of cell types in biology is widely debated,^38^ but the concept is useful as an heuristic device. This is where our spatial transcriptomics provides insight. It shows clearly that the cells that we define as CNT are in fact interposed between the DCT and the CD, harkening back to the original anatomical description of segment that connects DCT and CD. Yet, the robust and organized expression of transport and regulatory genes confirms Imai’s original suggestion that the CNT is a bona fide and unique ‘functional subdivision’ of the distal tubule.^2^

**Figure 8.**
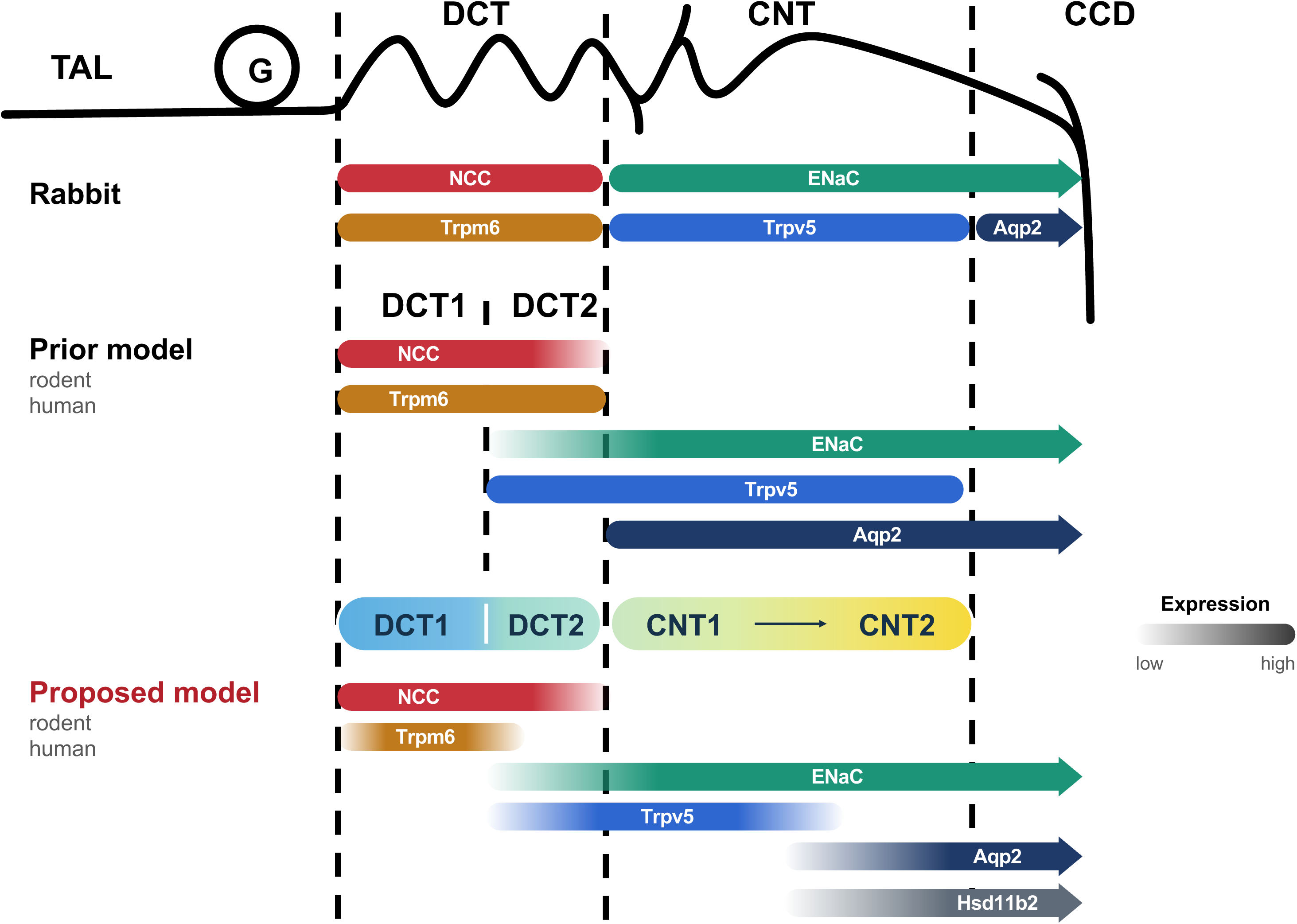
Proposed organization of calcium, sodium, and water transport pathways along the distal tubule. Distal tubule organization from Chen *et* al.,^9^ summarizing previously described species differences in distal tubule organization. The conventional rabbit model (upper) and prior rodent and human model (middle) are shown alongside the new model proposed from the findings of the present study (lower). The anatomical diagram indicates the transition from the TAL through the DCT and CNT to the CCD; dashed lines mark conventional segment boundaries. In the proposed model, DCT1 and DCT2 are followed by a gradual transition across CNT1 and CNT2, with intermediate transcriptional features between them. Colored bars depict the relative distributions of NCC, ENaC, TRPM6, TRPV5, AQP2, and 11β-hydroxysteroid dehydrogenase type 2 (HSD11B2). TRPV5-associated calcium transport machinery is concentrated toward DCT2 and CNT1, whereas AQP2 and HSD11B2 become prominent toward CNT2 and the CCD; ENaC-associated sodium transport machinery extends across the CNT. Color gradients indicate changes in relative expression. The bars are qualitative and do not represent measured transport rates or segment lengths. AQP2, aquaporin-2; CCD, cortical collecting duct; CNT, connecting tubule; DCT, distal convoluted tubule; ENaC, epithelial sodium channel; G, glomerulus; NCC, Na-Cl cotransporter; TAL, thick ascending limb; TRPM6, transient receptor potential cation channel subfamily M member 6; TRPV5, transient receptor potential cation channel subfamily V member 5.

---

**Supplemental Table 1:** Antibodies used in the study.

| Antibody | Host species | Supplier/source | Catalog number | IF dilution |
|---|---|---|---|---|
| **Total NCC** | Rabbit | D. H. Ellison laboratory | NA | 1:5000 |
| **AQP2** | Rabbit | Abcam | ab199975 | 1:3000 |
| **Parvalbumin** | Guinea pig | Swant | GP72 | 1:2000 |
| **GFP, FITC-conjugated** | Goat | Rockland | 600-102-215 | 1:500 |
| **Calbindin D28k (CALB1)** | Mouse | Swant | CB300 | 1:250 |
| **TRPV5** | Rabbit | Alomone | ACC-035 | 1:500 |

**Supplemental Table 2:** Plasma electrolytes Calbindin-INTACT mouse line.

| Plasma electrolytes | Mean ± SD (n = 7) |
|---|---|
| Na<sup>+</sup> (mmol/L) | 142.43 ± 2.37 |
| K<sup>+</sup> (mmol/L) | 3.81 ± 0.54 |
| Cl<sup>-</sup> (mmol/L) | 107.71 ± 7.48 |
| iCa<sup>2+</sup> (mmol/L) | 1.34 ± 0.07 |

