## Supplemental Figures for "Distinct calcium, sodium, and water programs characterize mouse and human connecting tubule"

#### Supplemental Figures, Legends, and References

##### Supplemental Figure Legends

**Supplemental Figure 1. GFP reporter labeling in relation to parvalbumin and calbindin-D28K expression in Calb1-Cre-INTACT kidneys.**
Representative overview of a mouse kidney cryosection from a Calb1-Cre-INTACT mouse immunostained for parvalbumin (red), GFP (green), and calbindin-D28K (white). Individual channels and the merged image are shown. Parvalbumin highlights DCT1 profiles, whereas calbindin-D28K marks overlapping distal nephron populations, including DCT2 and CNT. GFP labeling extends beyond profiles with detectable adult calbindin-D28K expression.
DCT, distal convoluted tubule; CNT, connecting tubule; GFP, green fluorescent protein.

**Supplemental Figure 2. GFP reporter labeling in relation to NCC and AQP2 expression in Calb1-Cre-INTACT kidneys.**
Representative kidney cryosection from a Calb1-Cre–INTACT mouse immunostained for aquaporin-2 (AQP2; red), GFP (green), and the Na–Cl cotransporter (NCC; white). Individual channels and the merged image shown. GFP, green fluorescent protein.

**Supplemental Figure 3. Cluster annotation and sample representation in the Calb1-Cre–INTACT snRNA-seq dataset.**
(A) Dot plot of canonical epithelial markers and selected transport-associated genes across the complete cluster annotations. The panel distinguishes DCT1 clusters, DCT2, CNT populations, collecting duct principal cells, ICA and ICB subclusters, urothelial populations, and proliferating DCT and CNT populations. (B) The shared UMAP embedding displayed separately for the two biological samples, a stacked bar showing the proportions of major annotated populations in the analyzed enriched dataset, and a dot plot of the canonical marker panel as described.[1] Both samples contribute to the major populations. Dot size represents the percentage of nuclei with detectable expression; color represents gene-wise scaled average expression. DCT, distal convoluted tubule; CNT, connecting tubule; PC-CD, collecting duct principal cells; PC-IMCD, inner medullary collecting duct principal cells; ICA and ICB, type A and type B intercalated cells; URO, urothelial cells; Prolif., proliferating cells; UMAP, uniform manifold approximation and projection.

**Supplemental Figure 4. Selected markers and differential expression profiles distinguish CNT populations.**
(A-D) Expression of *Slc8a1* (A), *Kl* (B), *Klk1* (C), and *Egfem1* (D) projected onto the UMAP of the Calb1-Cre-INTACT dataset. Each point represents one nucleus; purple intensity

indicates normalized expression, with a separate scale for each gene. (E) Dot plot of selected differentially expressed genes (DEG) across DCT1, DCT2, CNT1, $CNT_T$, CNT2, PC-CD, and the remaining annotated populations grouped as “Others.” Note that PC-CD cells does not included PC-IMCD. Markers were identified using Seurat’s FindAllMarkers function with a Wilcoxon rank-sum test, retaining positively enriched genes with a minimum log2 fold change of 0.5 and detection in at least 25% of nuclei in either comparison group. P values were adjusted using the Bonferroni method. Dot size indicates the percentage of expressing nuclei, and color indicates gene-wise scaled average expression. The profiles show shared and population-associated expression across the three CNT groups. Complete DEG list is in supplemental material. CNT, connecting tubule; $CNT_T$, transitional CNT; DCT, distal convoluted tubule; PC-CD, collecting duct principal cells; UMAP, uniform manifold approximation and projection.

**Supplemental Figure 5. Calcium, sodium & potassium, and water gene modules across distal epithelial populations.**
UMAP projections of module scores for calcium transport (upper left), water transport (upper right), and sodium & potassium transport (lower left) in the Calb1-Cre–INTACT dataset. Calcium-module genes were *Slc8a1, Calb1, Trpv5, S100g, Kl, Vdr, Pth1r*, and *Casr*; water-module genes were A*qp2, Aqp3, Aqp4*, and *Avpr2*; and sodium/potassium-module genes were *Scnn1a, Scnn1b, Scnn1g, Hsd11b2*, *Sgk1, Nr3c2*, and *Kcnj1*. Scores were calculated using Seurat’s AddModuleScore function relative to expression-matched control genes. Each point represents one nucleus, and color denotes the module score, with display limits set to the 5th and 95th percentiles for each module. Calcium-associated scores are prominent in DCT2 and CNT1, water-associated scores toward CNT2 and collecting duct populations, and sodium/potassium-associated scores across CNT populations. CNT, connecting tubule; DCT, distal convoluted tubule; UMAP, uniform manifold approximation and projection.

**Supplemental Figure 6. Canonical markers and sodium transport genes across major distal tubule cell types.** (A) UMAP of the Calb1-Cre–INTACT dataset showing all major distal tubule cell types. (B) Dot plots of canonical cell-type markers (left) and genes involved in sodium transport and its regulation (right) across the annotated populations. Dot size indicates the percentage of nuclei expressing each gene; color indicates gene-wise scaled average expression. DCT, distal convoluted tubule; CNT, connecting tubule; PC-CD, collecting duct principal cells; PC-IMCD, inner medullary collecting duct principal cells; ICA, type A intercalated cell; ICB, type B intercalated cell; URO, urothelium; Prolif, proliferating cells. UMAP, uniform manifold approximation and projection.

**Supplemental Figure 7. Expression of GPCR-related, epithelial identity and transcription factor genes across major distal tubule cell populations.** Dot plots show selected genes related to G protein-coupled receptor signaling (upper left), epithelial identity, ciliary function, and acid–base transport (upper right), and transcriptional regulation (lower left) across the major populations in the Calb1-Cre–INTACT dataset. Dot size indicates the percentage of nuclei with detectable expression, and color indicates gene-wise scaled average expression CNT, connecting tubule; $CNT_T$, transitional CNT; DCT, distal convoluted tubule; PC-CD, collecting duct principal cells; PC-IMCD, inner medullary collecting duct principal cells; ICA and ICB, type A and type B intercalated cells; URO, urothelial cells; Prolif., proliferating cells.

**Supplemental Figure 8. Nephron- and ureteric-associated marker expression across CNT populations.**
(A) UMAP feature plots of the nephron-associated markers *Slc2a9* and *Hoxd10*. (B) Expression of the ureteric-associated marker *Ache*. (C) Umap displaying all annotated populations. (D) Dot plot of selected lineage-associated and segment-specific markers across the annotated populations. Markers were drawn from Ransick et al.[2] Feature-plot color indicates normalized expression, with gene-specific scales. In the dot plot, size indicates the percentage of expressing nuclei and color indicates gene-wise scaled average expression. *Slc2a9* expression is prominent in DCT2, CNT1, and $CNT_T$, whereas detectable *Ache* expression associates CNT2 with collecting duct populations. CNT, connecting tubule; $CNT_T$, transitional CNT; DCT, distal convoluted tubule; PC-CD, collecting duct principal cells; PC-IMCD, inner medullary collecting duct principal cells; ICA and ICB, type A and type B intercalated cells; URO, urothelial cells; Prolif., proliferating cells; UMAP, uniform manifold approximation and projection.

**Supplemental Figure 9. Comparison with published segment-specific markers from Ransick et al. within the Calb1-Cre–INTACT dataset.** (Left) Umap displaying all Calb1-Cre–INTACT dataset annotated populations. (Right) Dot plot of segment-specific marker genes defined by Ransick et al.[2] across annotated cell types in the d Calb1-Cre–INTACT dataset. Dot size indicates the percentage of nuclei expressing each gene, and color indicates scaled average expression. Marker genes are grouped by their original annotation in Ransick et al., and vertical lines mark the cell types expected to express each group. CCD, cortical collecting duct; OMCD, outer medullary collecting duct; IMCD, inner medullary collecting duct; dME, deep medullary epithelium; CNT, connecting tubule; $CNT_T$, transitional CNT; DCT, distal convoluted tubule; ICA and ICB, type A and type B intercalated cells; URO, urothelial cells; Prolif., proliferating cells; UMAP, uniform manifold approximation and projection.

**Supplemental Figure 10. (A)** Heatmap of 351 reference pairwise marker genes across CNT1, $CNT_T$ and CNT2 in the snRNA-seq reference and two Xenium replicates. Values are mean log2 expression, rows are z-scored within each dataset. Rows are grouped by the state with the highest snRNA-seq expression: CNT1-high (n = 157), $CNT_T$-high (n = 56) and CNT2-high (n = 138). **(B)** Concordance of log2 fold changes between Xenium and snRNA-seq for each pairwise comparison (CNT1 vs $CNT_T$, n = 182; $CNT_T$ vs CNT2, n = 175; CNT1 vs CNT2, n = 324) and each replicate. Metrics are Spearman ρ, the percentage of markers with concordant direction, and the regression slope of Xenium on snRNA-seq log2 fold change.

**Supplemental Figure 11. Program-level concordance between snRNA-seq and Xenium CNT classifications.** UMAPs and box plots show CNT nuclei from the snRNA-seq reference (left) and CNT cells from Xenium replicates 1 (middle) and 2 (right). Programs were derived by cNMF (K = 6) of snRNA-seq CNT nuclei restricted to Xenium panel genes and projected onto Xenium cells with starCAT. **(A)** CNT state annotation. **(B–D)** Usage of the CNT1-like (B), $CNT_T$-like (C) and CNT2-like (D) programs, normalized across the three programs. **(E)** CNT1-to-CNT2 program score, defined as CNT2-like minus CNT1-like usage. **(F)** CNT1-to-CNT2 program score by annotated state. Boxes indicate median and interquartile range; whiskers extend to 1.5 × interquartile range.

**Supplemental Figure 12. Regional distribution of distal tubule epithelial populations in Xenium sample 1.**

Publicly available Xenium data from male mouse kidney[3] were reanalyzed to identify populations corresponding to those in the Calb1-Cre–INTACT snRNA-seq dataset. (A) UMAP showing the annotated distal tubule populations within sample 1; remaining kidney populations are gray. (B) Tissue map colored by anatomical region: cortex, outer stripe of the outer medulla (OSOM), inner stripe of the outer medulla (ISOM), inner medulla, and adjacent fat as delineated in (Bahena-Lopez et al.). The corticomedullary distance convention is indicated below the map: zero denotes the outer-inner medullary boundary, negative distances extend toward the cortex, and positive distances extend into the inner medulla. (C) Dot plot of Xenium dataset sample 1 showing key markers. Dot size indicates the percentage of cells expressing each gene, and color indicates gene-wise scaled average expression. (D) The same UMAP displayed by the anatomical region. This figure presents sample 1 of two biological replicates. CNT, connecting tubule; $CNT_T$, transitional CNT; DCT, distal convoluted tubule; PC-CD, collecting duct principal cells; ICA and ICB, type A and type

B intercalated cells; URO, urothelial cells; snRNA-seq, single-nucleus RNA sequencing; UMAP, uniform manifold approximation and projection.

**Supplemental Figure 13. Regional distribution of distal tubule epithelial populations in Xenium sample 2.**
Publicly available Xenium data from male mouse kidney[3] were reanalyzed to identify populations corresponding to those in the Calb1-Cre–INTACT snRNA-seq dataset. (A) Spatial map of sample 2, with annotated distal tubule populations colored as indicated and other kidney cell types shown in gray. (B) The outlined cortical region is enlarged in B. The same region displayed without (left) and with (right) intercalated cells highlighted, showing the arrangement of DCT and CNT populations within tubular profiles. (C) Dot plot of Xenium dataset sample 2 showing key markers. Dot size indicates the percentage of cells expressing each gene, and color indicates gene-wise scaled average expression. (D) Tissue map colored by anatomical region: cortex, outer stripe of the outer medulla (OSOM), inner stripe of the outer medulla (ISOM), inner medulla, and adjacent fat as delineated in[3]. The corticomedullary distance convention is indicated below the map: zero denotes the outer-inner medullary boundary, negative distances extend toward the cortex, and positive distances extend into the inner medulla. (E) UMAP showing the annotated distal tubule populations within sample 2 by the anatomical region. This figure presents sample 2 of two biological replicates. CNT, connecting tubule; $CNT_T$, transitional CNT; DCT, distal convoluted tubule; PC-CD, collecting duct principal cells; ICA and ICB, type A and type B intercalated cells; URO, urothelial cells; snRNA-seq, single-nucleus RNA sequencing; UMAP, uniform manifold approximation and projection.

**Supplemental Figure 14.** Heterotypic spatial adjacency of distal tubule cell types in Xenium spatial transcriptomics. (A) Heterotypic adjacency among cortical distal tubule cell types (DCT2, CNT1, $CNT_T$, CNT2 and PC-CD), shown separately for each Xenium biological replicate. For each source cell type (rows), the heatmap gives the fraction of neighboring cells of a different type, within a 20 μm radius, that belong to each neighbor cell type (columns). Same-type neighbors were excluded, and each row sums to 100%. (B) Heterotypic adjacency among all distal tubule cell populations, combining two biological replicates. DCT, distal convoluted tubule; CNT, connecting tubule; PC-CD, principal cells of the collecting duct; PC, principal cell; ICA, type A intercalated cell; ICB, type B intercalated cell; URO, urothelium.

**Supplemental Figure 15. Pseudospatial arrangement of calcium, sodium and water transport gene expression from DCT2 to the PC-CD in cortical regions.** Pseudotime-ordered heatmaps incorporating both Xenium samples, restricted to the cortical DCT2–

CNT–PC-CD populations and rooted in DCT2. Smoothed expression of calcium-, sodium-, and water-associated genes is displayed as gene-wise z-scores, with cell-type annotations above.

**Supplemental Figure 16. Corticomedullary expression profiles distinguish calcium- and water-associated genes and the distribution of principal and intercalated cells in Xenium spatial transcriptomics.** (A) Gene expression and cell distributions were evaluated along the anatomically defined corticomedullary distance axis used in the original Xenium study[3]. Zero marks the boundary between the outer and inner medulla; negative distances extend through the outer medulla toward the cortex, and positive distances extend into the inner medulla. Distances are expressed in micrometers and represent tissue position, not distance traveled along a tubule. Values summarize two male mouse kidneys. Mean SCT-normalized expression of *Avpr2*, *Aqp2*, *Calb1*, and *Trpv5* within 250-μm distance bins, calculated separately for each sample and then averaged across both biological replicates. *Calb1* and *Trpv5* are most prominent at cortical positions and decline toward the OSOM, whereas *Aqp2* and *Avpr2* increase toward medullary positions and remain elevated in the inner medulla. The dashed vertical line marks zero. These profiles recapitulate the regional distinction between calcium- and water-associated expression identified in the Calb1-Cre–INTACT snRNA-seq dataset. (B) Mean numbers of type A intercalated cells (ICA), type B intercalated cells (ICB), and principal cells (PC) in 100-μm distance bins across the two samples; red vertical lines mark zero. ICB are concentrated at cortical positions, ICA extend across cortical and medullary positions, and PC counts are greatest in the inner medulla. Counts are not normalized to tissue area and should be interpreted in relation to the sampled tissue coverage.
SCT, SCTransform; snRNA-seq, single-nucleus RNA sequencing.

**Supplemental Figure 17. Gaussian mixture model identification of human $CNT_T$. (A)** Human CNT nuclei (n = 12,559) plotted by the CNT1-end and CNT2-end program scores, colored by Gaussian mixture model assignment. Each score is scaled so that 1 equals the mean level at the corresponding end of the CNT1-to-CNT2 axis. Crosses mark component means. The dashed box marks the region where both scores lie between 0.25 and 0.75. Nuclei with a grouped posterior probability below 0.8 are uncertain. **(B)** Mixture model assignments on the human CNT UMAP: CNT1-like (29.0%), both ($CNT_T$; 16.5%), CNT2-like (35.4%) and uncertain (19.1%). **(C)** Human $CNT_T$ nuclei highlighted on the UMAP. **(D)** Positive control test in which the same Gaussian mixture model was applied to CNT nuclei from our distal-enriched mouse snRNA-seq dataset (n = 6,892). **(E)** Mouse mixture model assignments versus the original annotation, shown as row percentages.

**Supplemental Figure 18. Direct projection of mouse CNT labels from distal-enriched snRNA-seq to human and mouse whole-kidney snRNA-seq. (A–C)** Human CNT nuclei from the KPMP Atlas v2 whole-kidney snRNA-seq dataset (n = 12,559), with mouse CNT labels transferred using 10 (A), 20 (B) and 30 (C) dimensions. Prediction scores were averaged over 30 nearest neighbors, and each nucleus was assigned the state with the highest score. **(D–F)** Usage of the CNT1-like, $CNT_T$-like and CNT2-like programs from full-gene cNMF (K = 5) of mouse distal-enriched snRNA-seq, projected onto 12,559 human CNT nuclei from the KPMP Atlas v2 whole-kidney snRNA-seq dataset using starCAT with 1,206 one-to-one mouse–human orthologs. **(G-I)** CNT nuclei from the KPMP mouse whole-kidney snRNA-seq dataset (n = 2,149 from 31 mice), shown by original atlas subclass label (G) and by mouse CNT labels transferred using 10 dimensions, before (H) and after (I) nearest-neighbor smoothing.

**Supplemental Figure 19. Gene-level and single-cell evidence for an intermediate CNT state between CNT1 and CNT2 in five datasets.** $CNT_T$ groups were defined differently in each dataset. In mouse distal-enriched snRNA-seq and Xenium, they are the original annotations. In mouse whole-kidney snRNA-seq, they are labels transferred from the distal-enriched reference. In human whole-kidney snRNA-seq, they are Gaussian mixture model assignments, with uncertain nuclei excluded. CNT1 and CNT2 cells were randomly split in half: half A was used to select CNT1-versus-CNT2 differentially expressed genes, and half B to measure expression levels. **(A)** For each CNT1-versus-CNT2 differentially expressed gene, mean expression in $CNT_T$ relative to CNT1 (y-axis) is plotted against mean expression in CNT2 relative to CNT1 (x-axis), both as log2 differences. Dashed lines mark $CNT_T$ expression equal to CNT1 (blue) and equal to CNT2 (orange). Genes are classified by whether $CNT_T$ expression lies between those of CNT1 and CNT2, above both or below both. **(B)** Single-cell CNT1-gene and CNT2-gene scores, each scaled to the median levels of half-B CNT1 and CNT2 cells, for $CNT_T$ cells and half-B CNT1 and CNT2 cells. Dotted lines mark scores of 0.25, and the shaded region marks cells with both scores above 0.25. Black contours show the two-dimensional density of $CNT_T$ cells.

Supplemental Figure 1

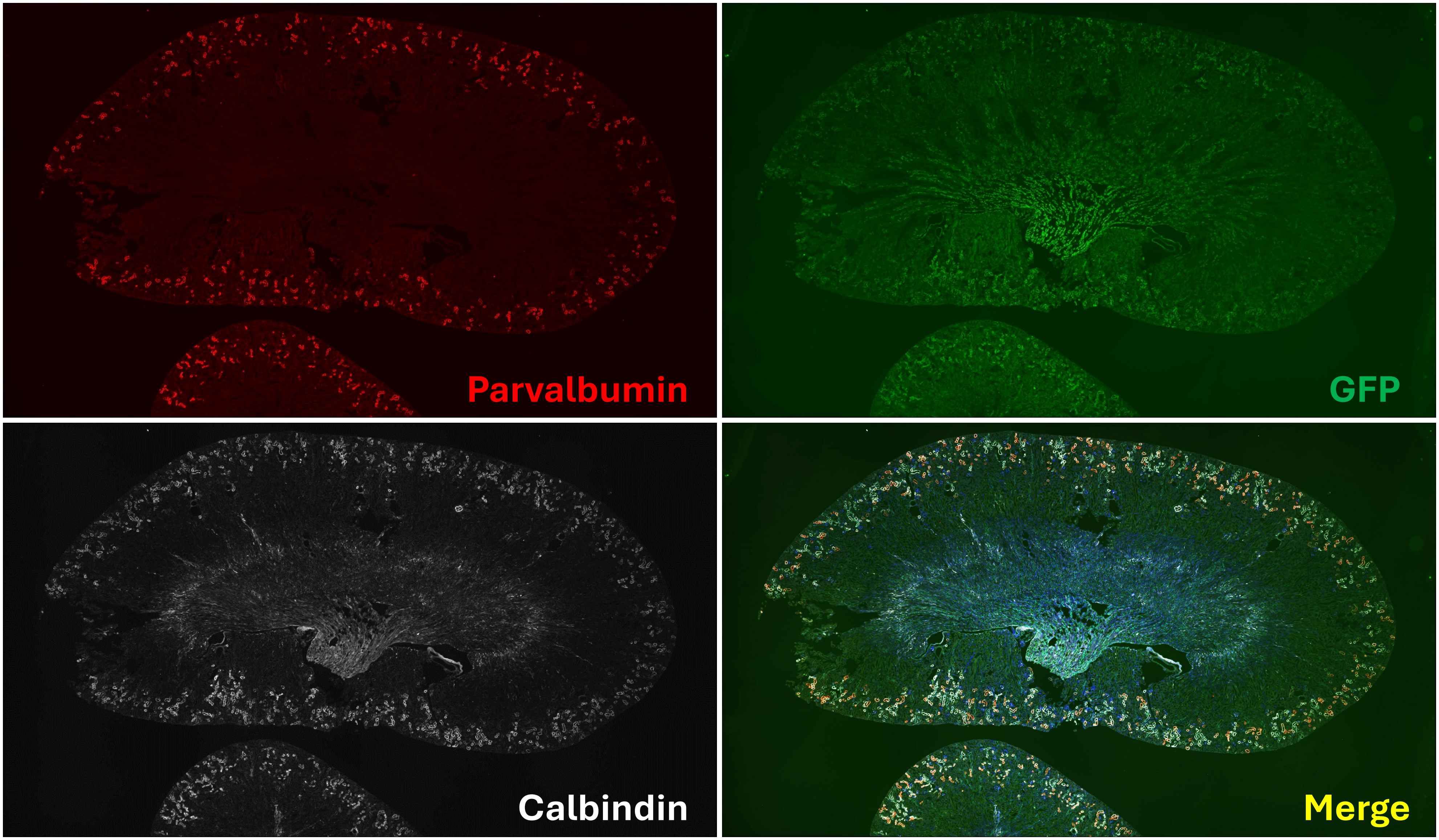

Supplemental Figure 2

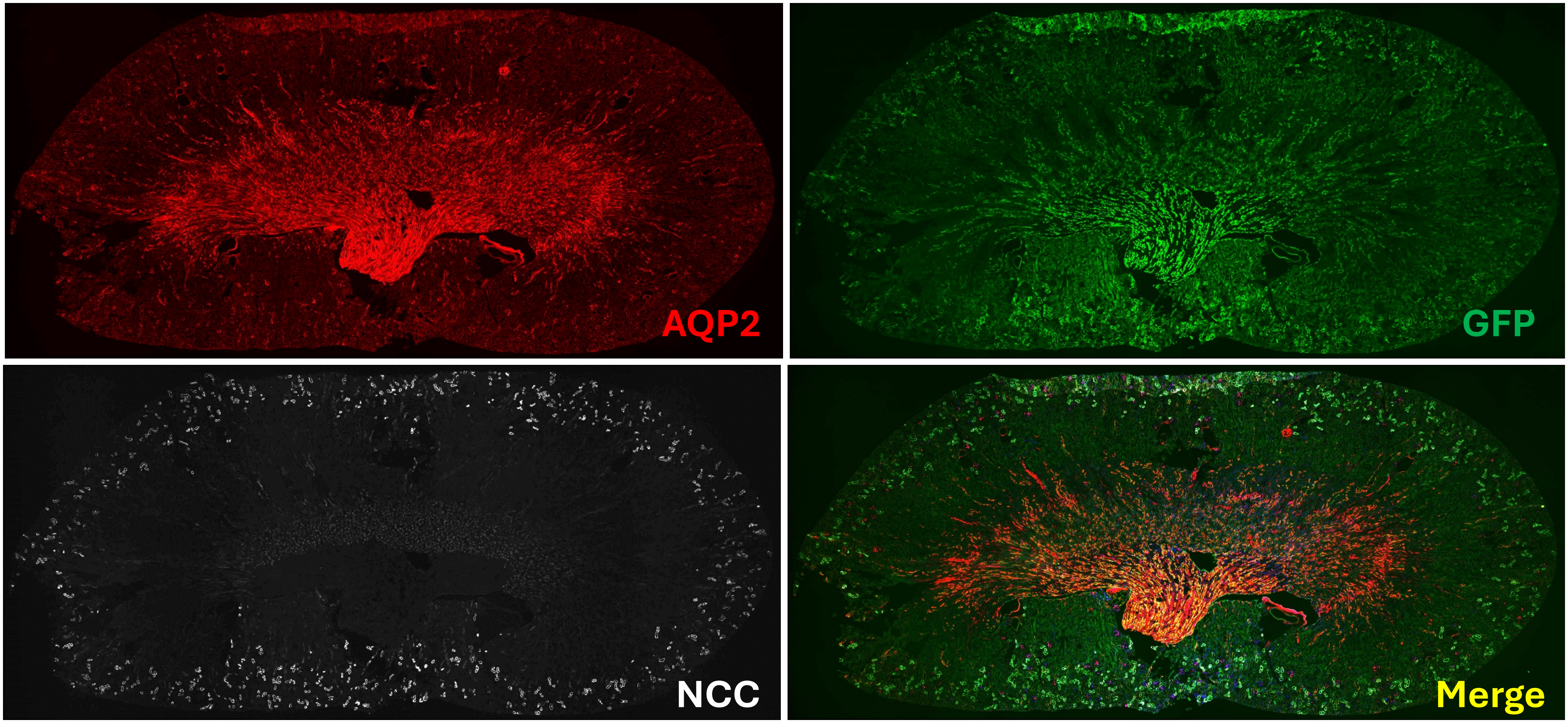

Supplemental Figure 3

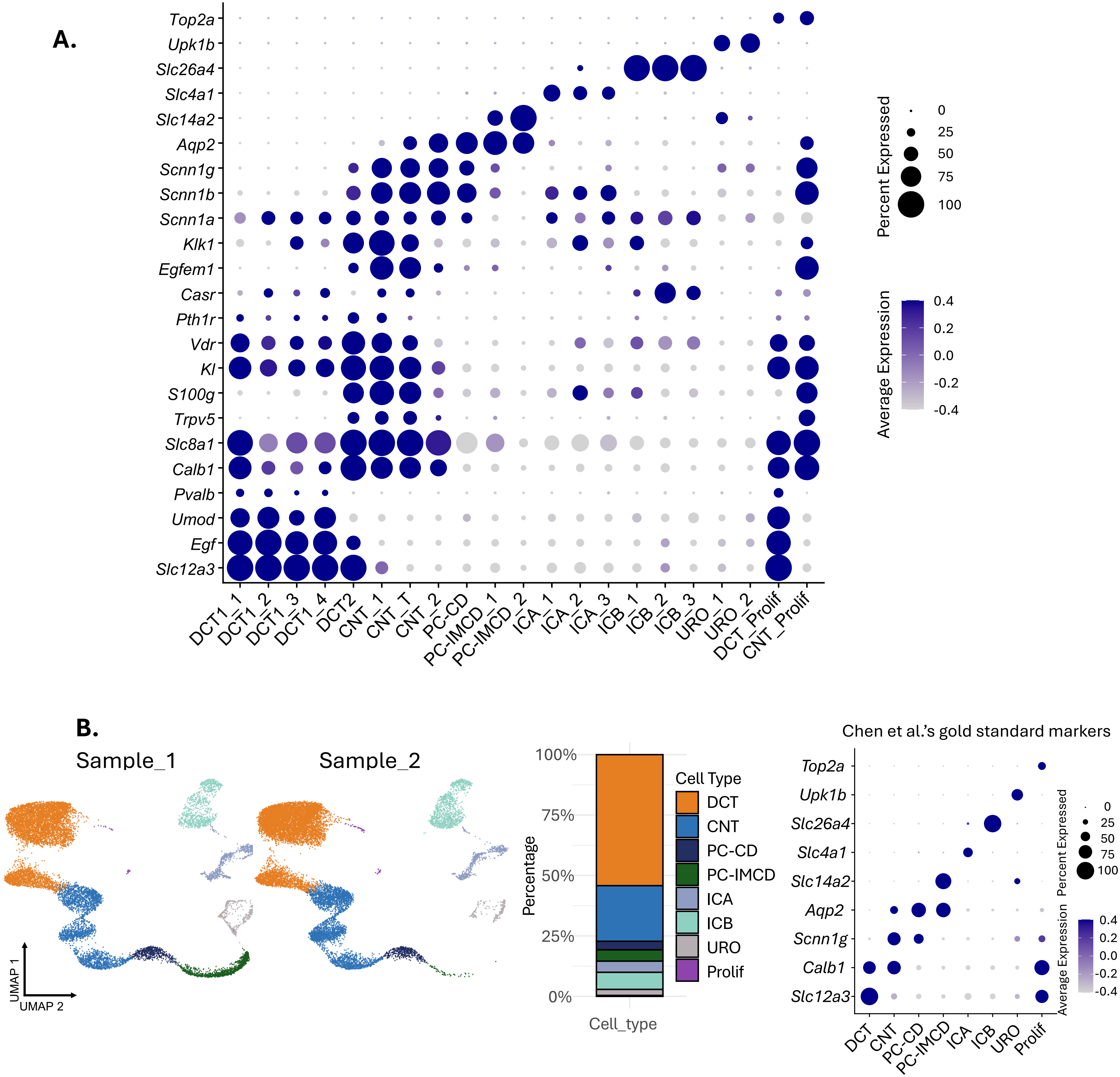

Supplemental Figure 4

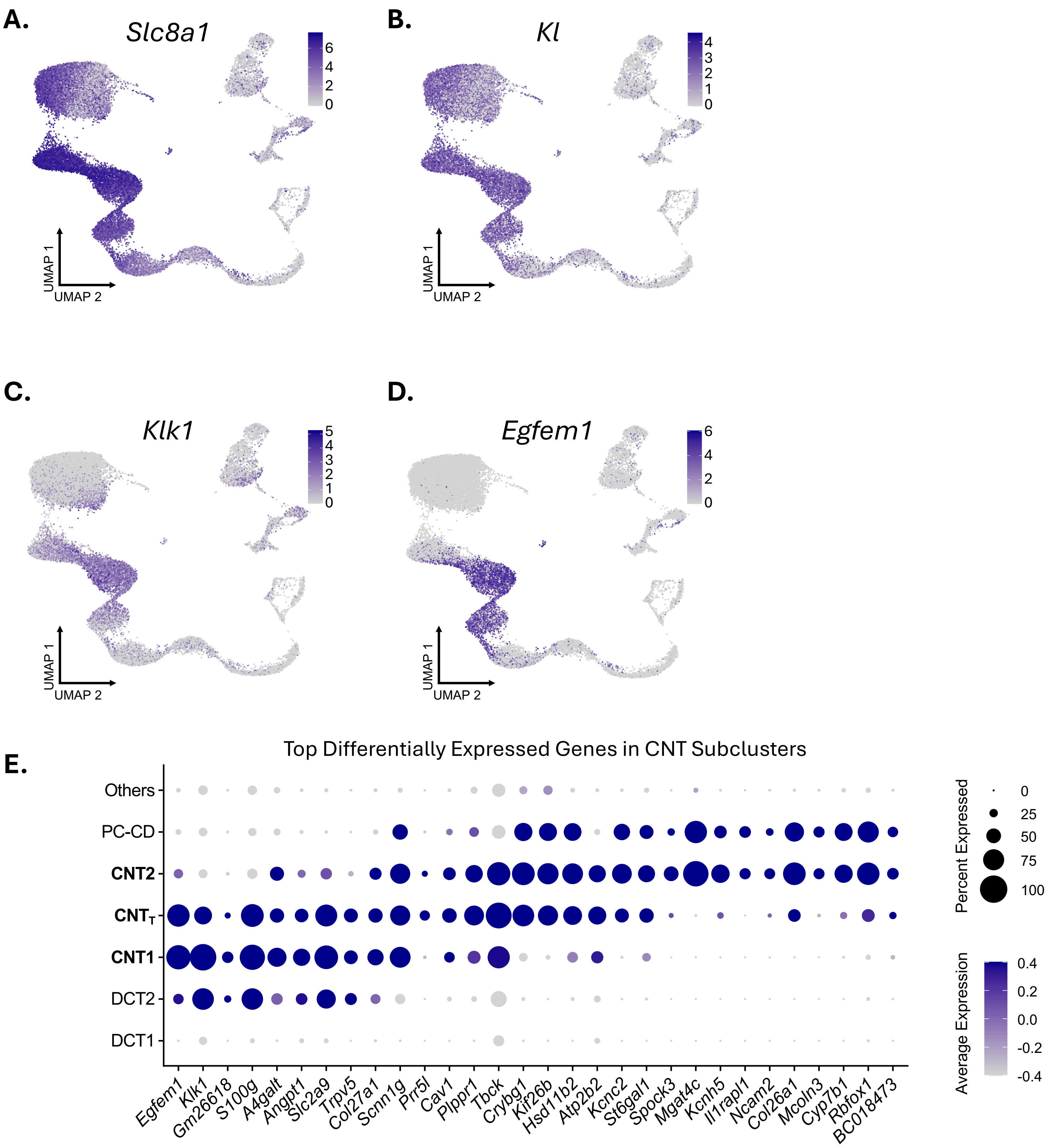

Supplemental Figure 5

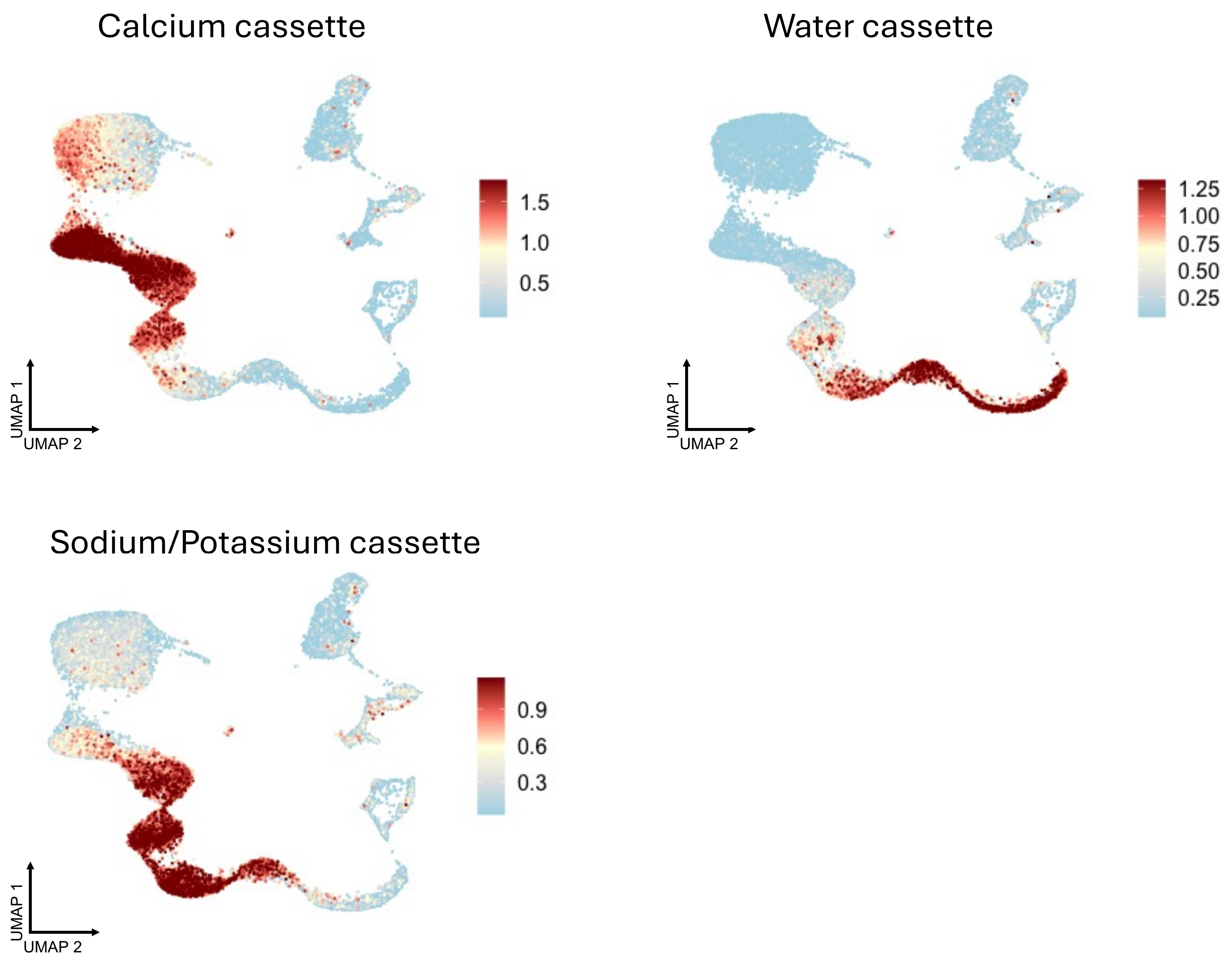


- Calcium cassette: *Slc8a1, Calb1, Trpv5, S100g, Kl, Vdr, Pth1r, Casr*.
- Water cassette : *Aqp2, Aqp3, Aqp4, Avpr2*.
- Sodium and Potassium cassette CNT: *Scnn1a, Scnn1b, Scnn1g, Hsd11b2, Sgk1, Nr3c2, Kcnj1*.

Supplemental Figure 6

A.

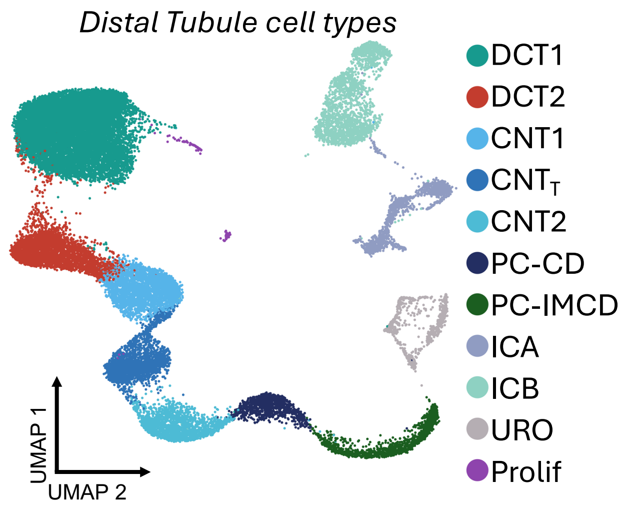


B.

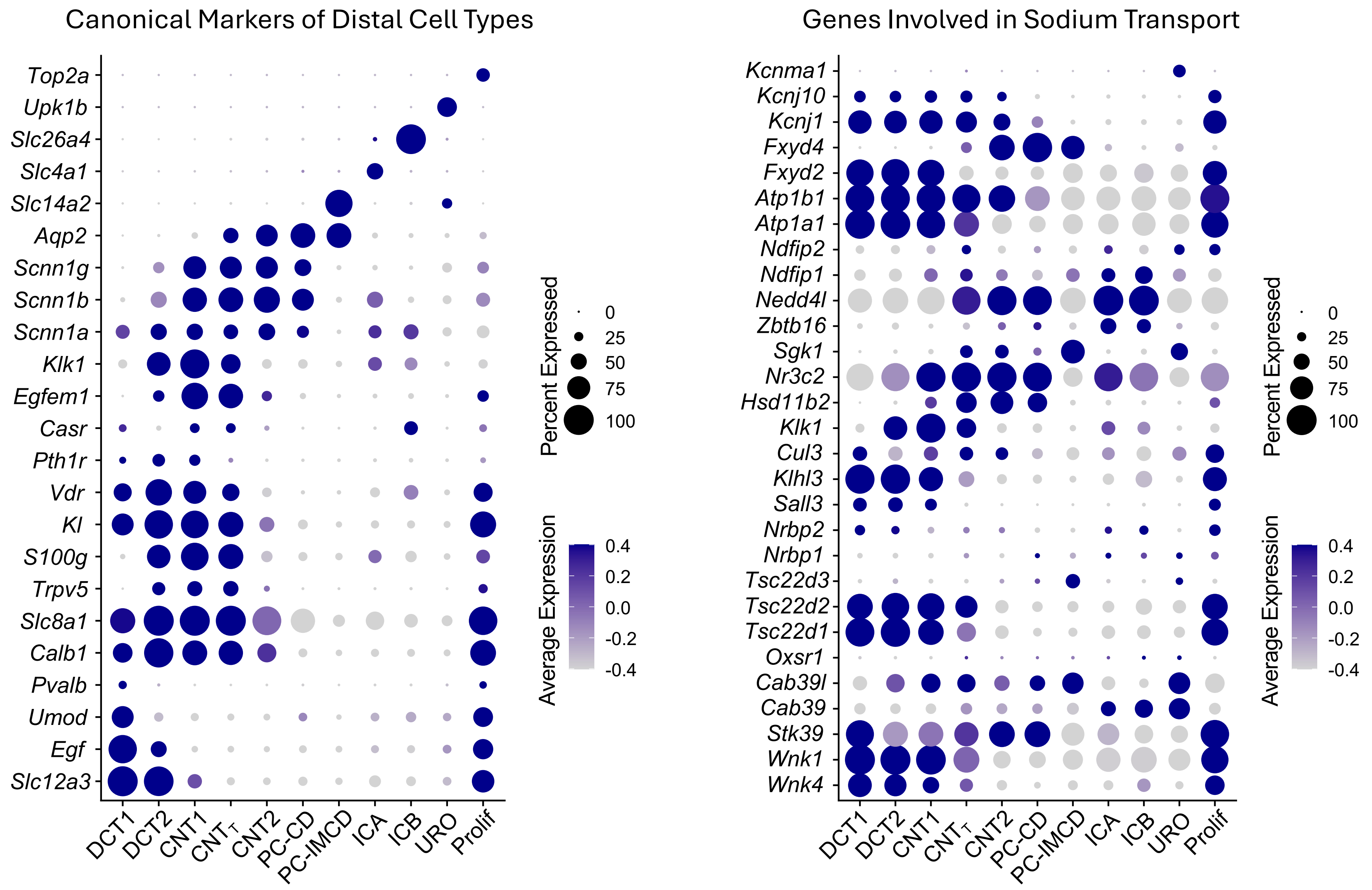

#### Supplemental Figure 7

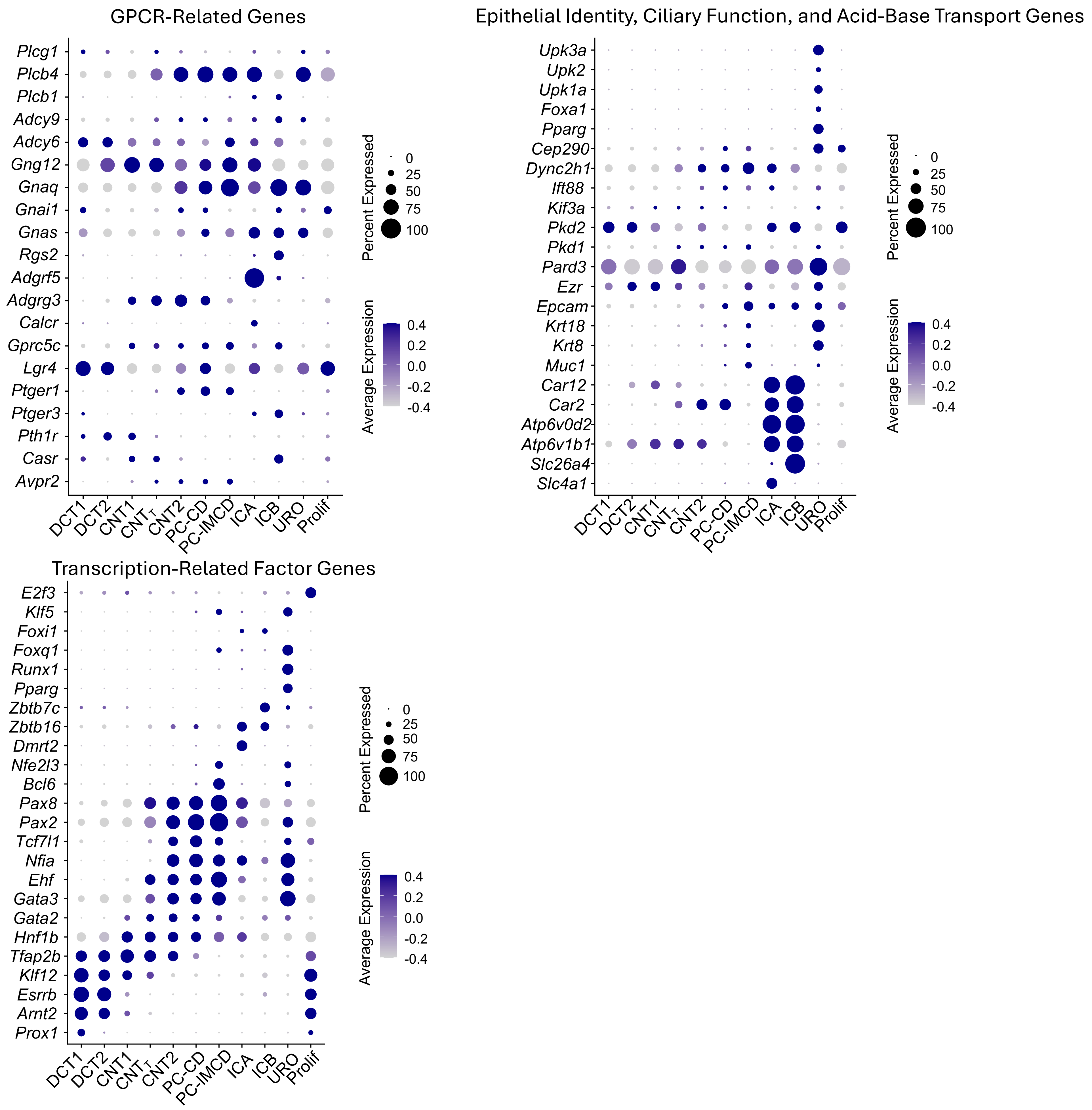

Supplemental Figure 8

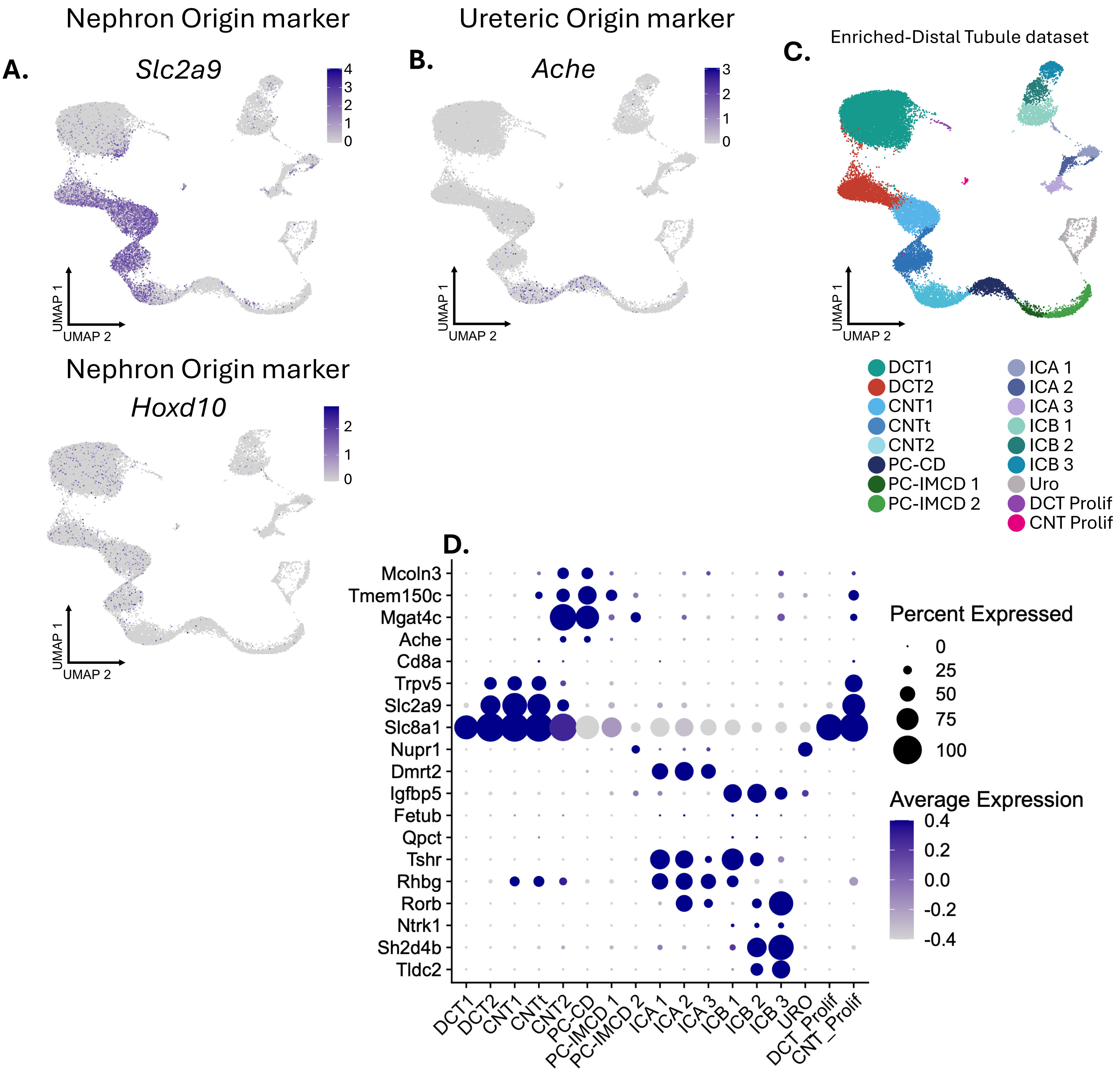

### Supplemental Figure 9

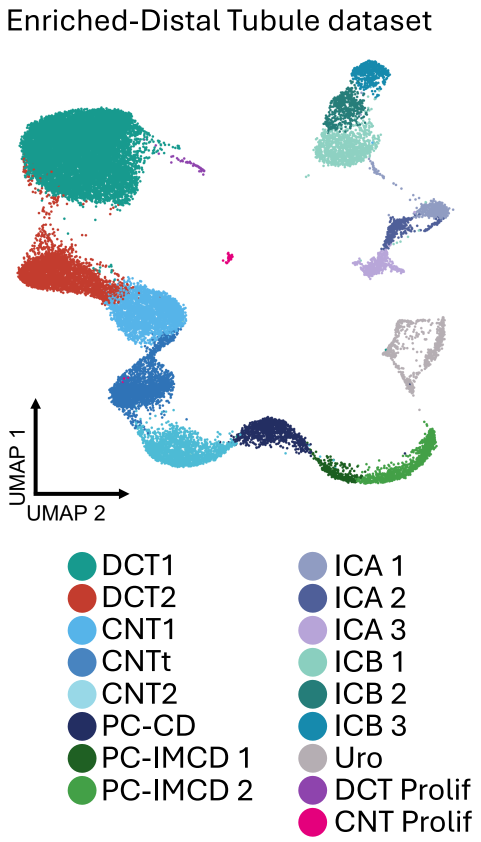


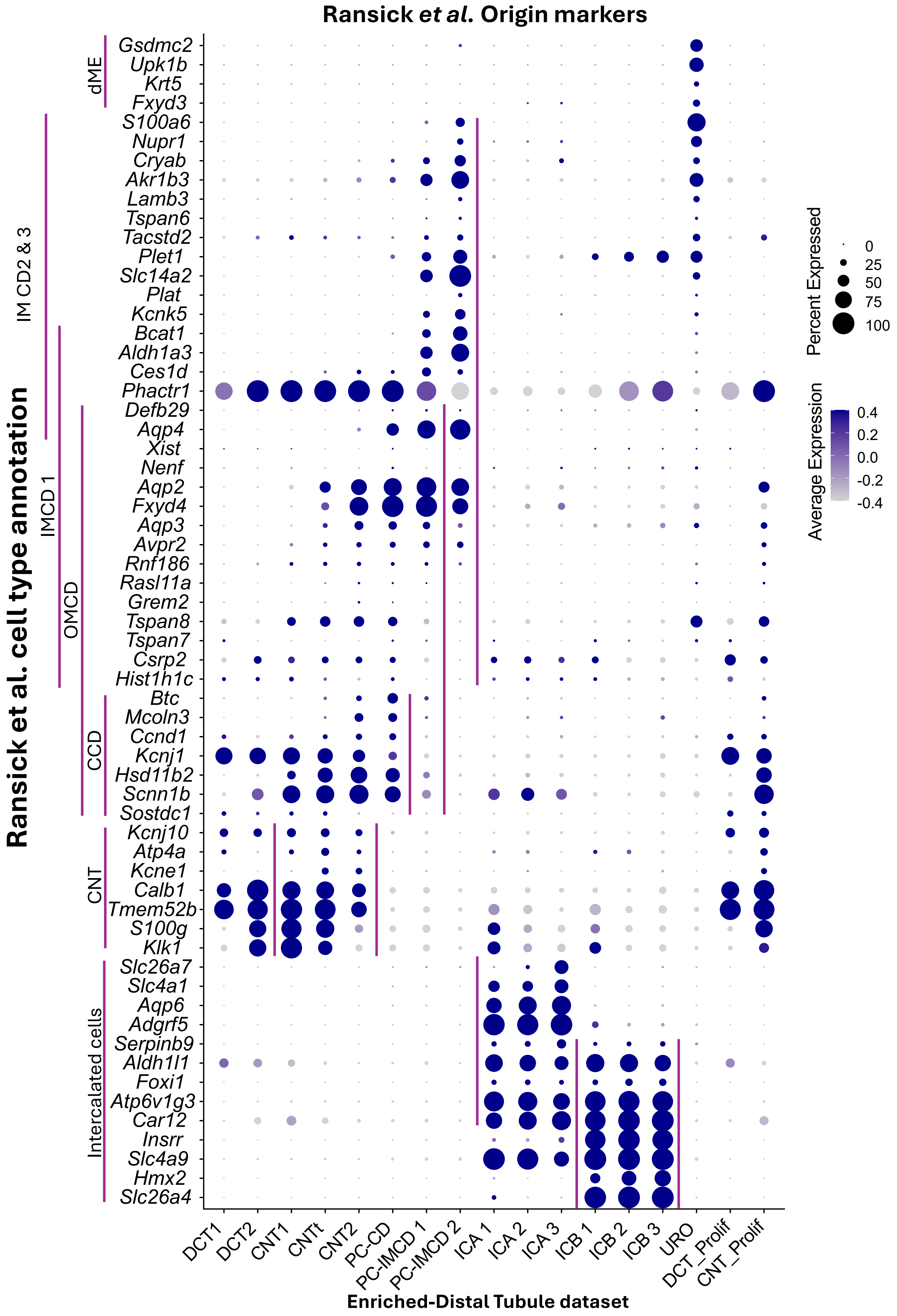

Supplemental Figure 10

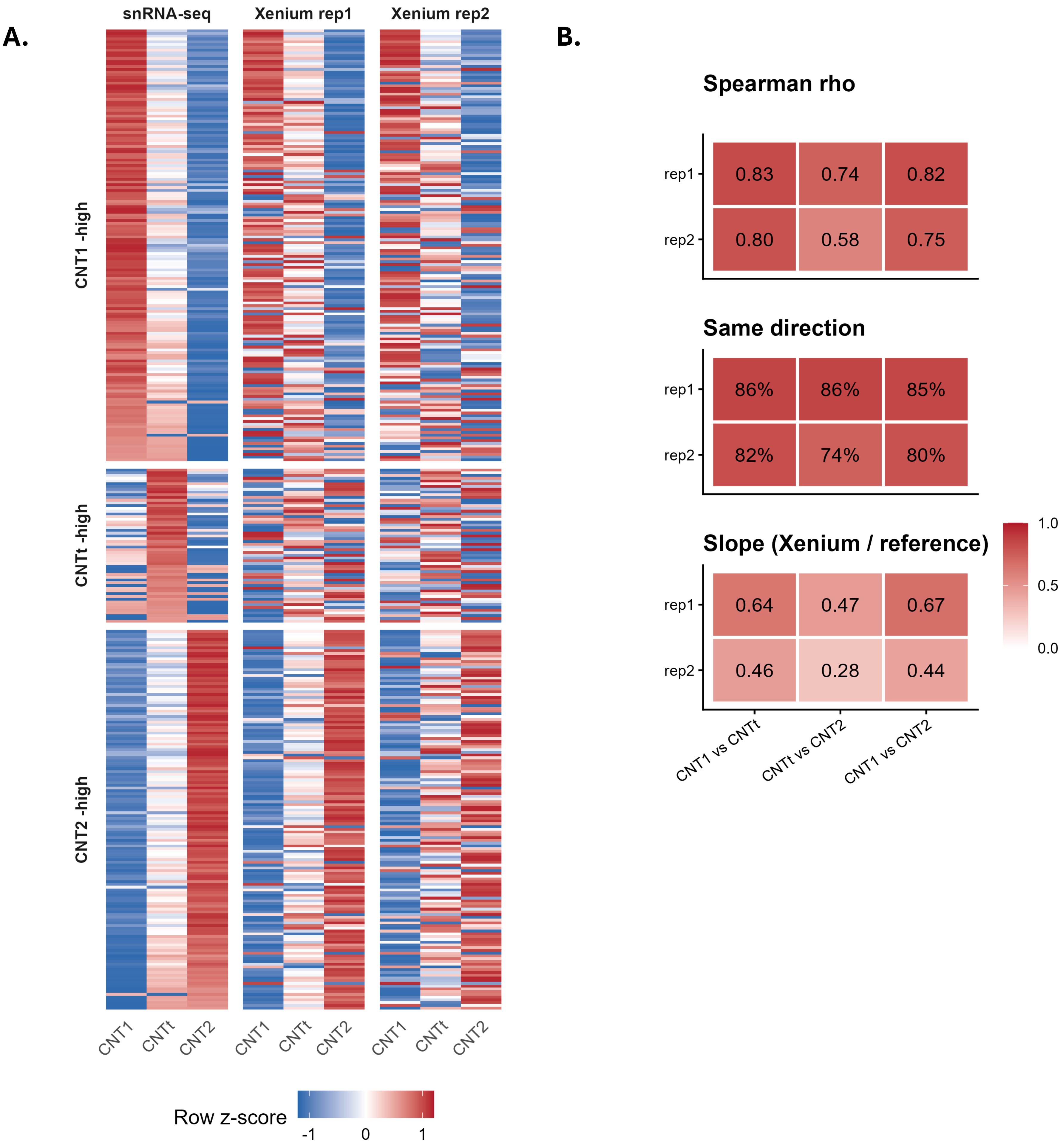

Supplemental Figure 11

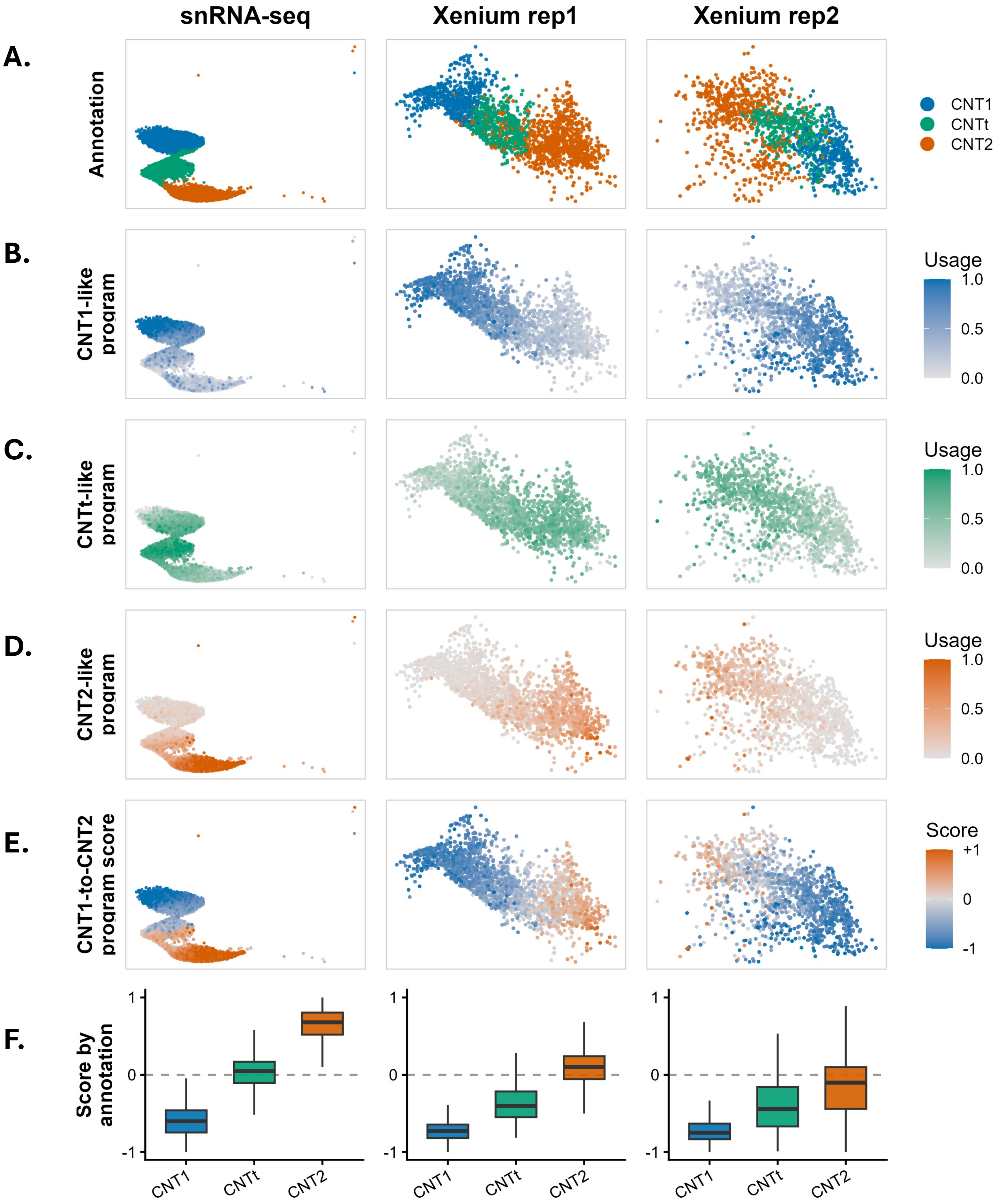

#### Supplemental Figure 12

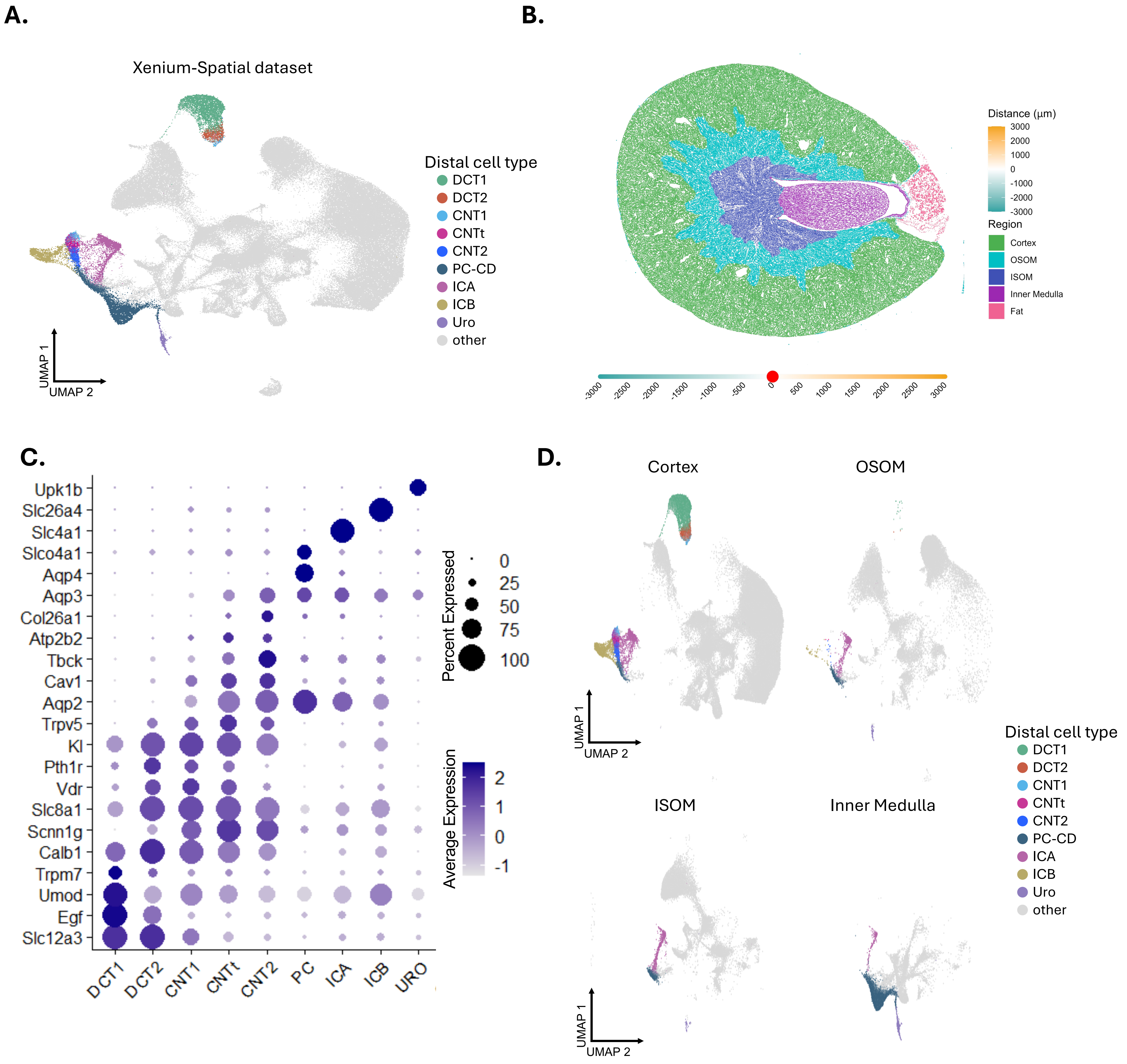

### Supplemental Figure 13

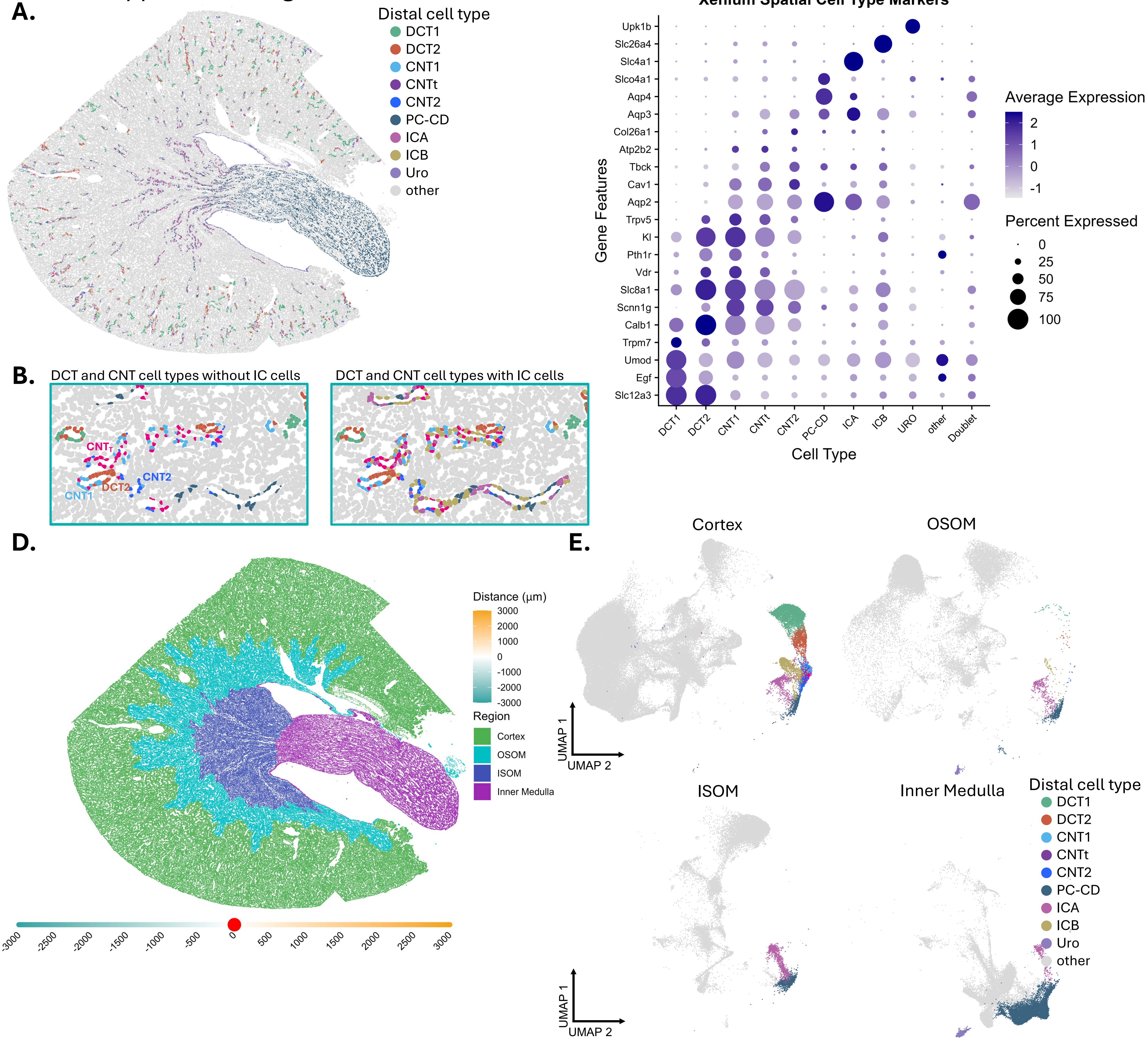

Supplemental Figure 14

A.

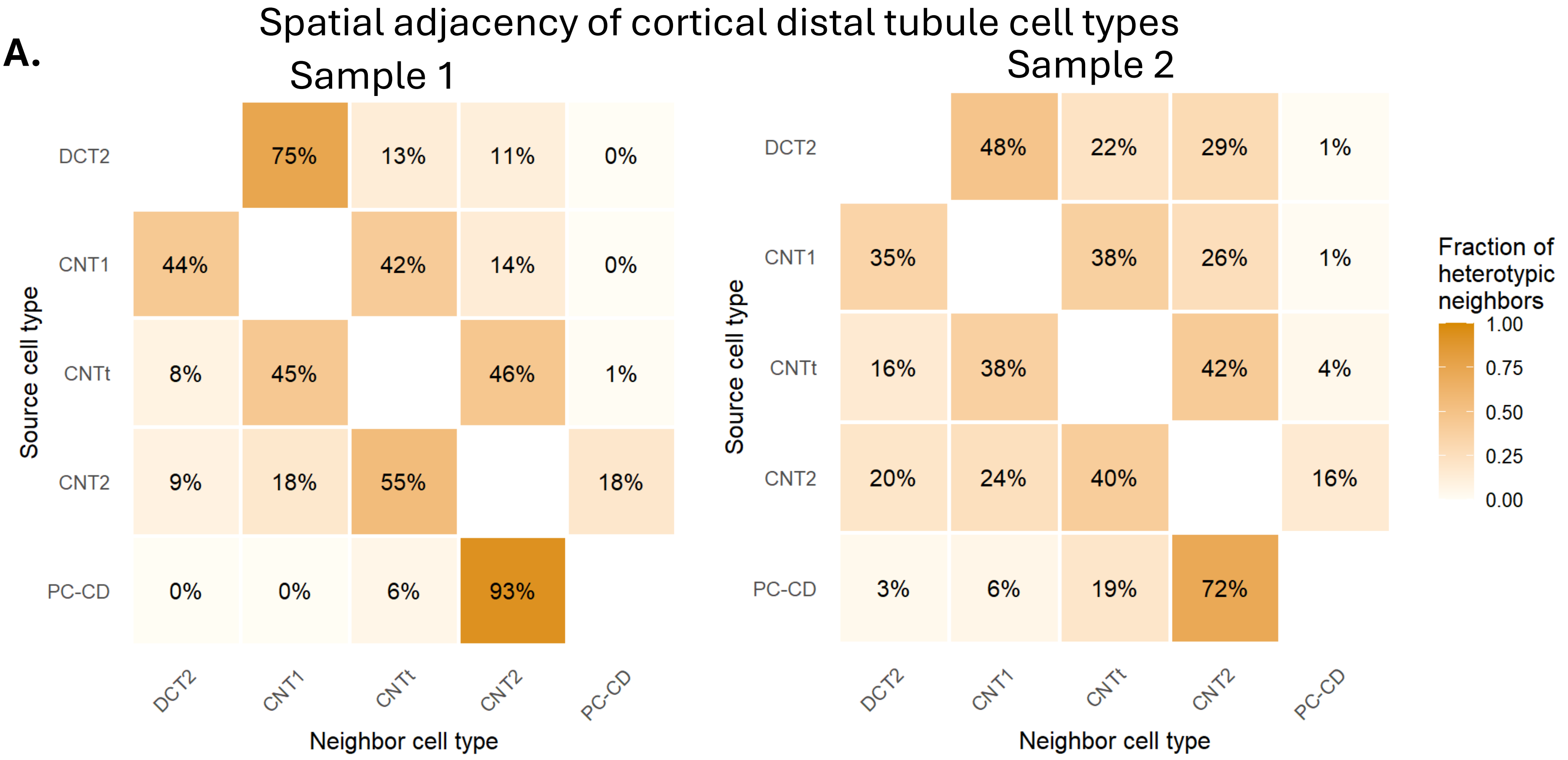


B.

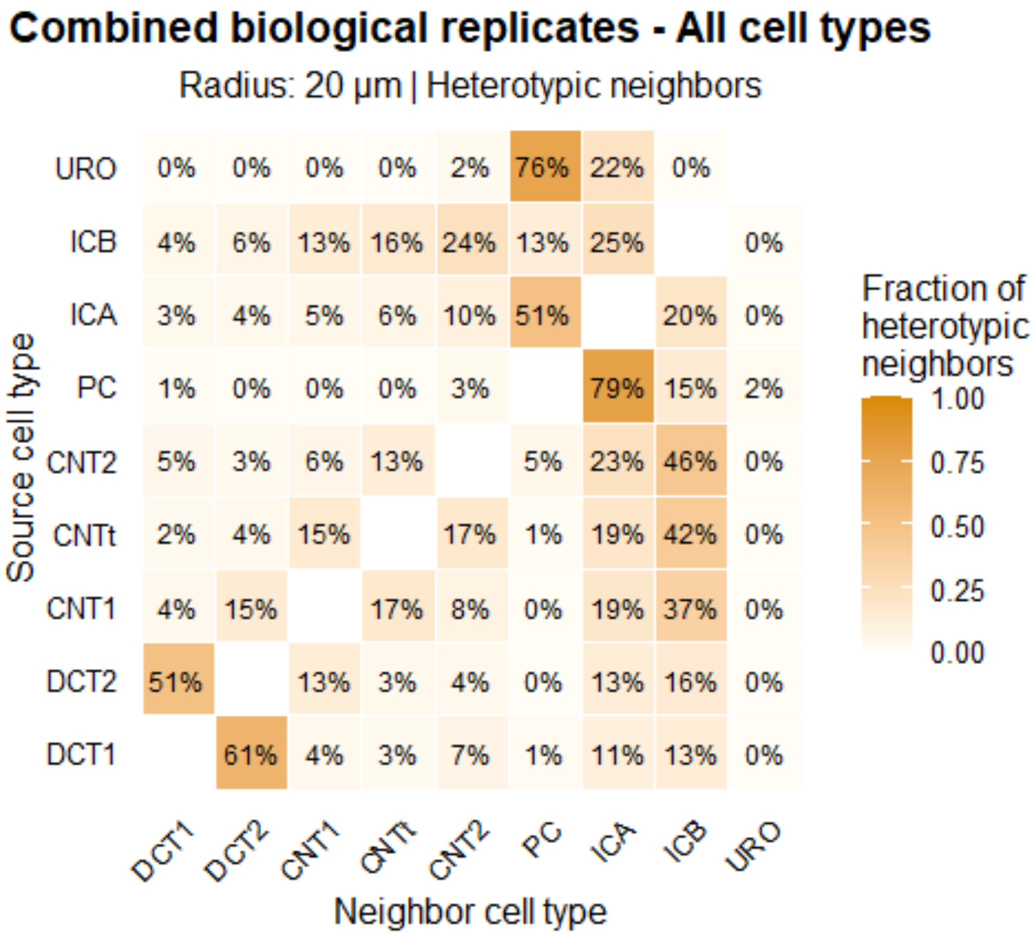

Supplemental Figure 15

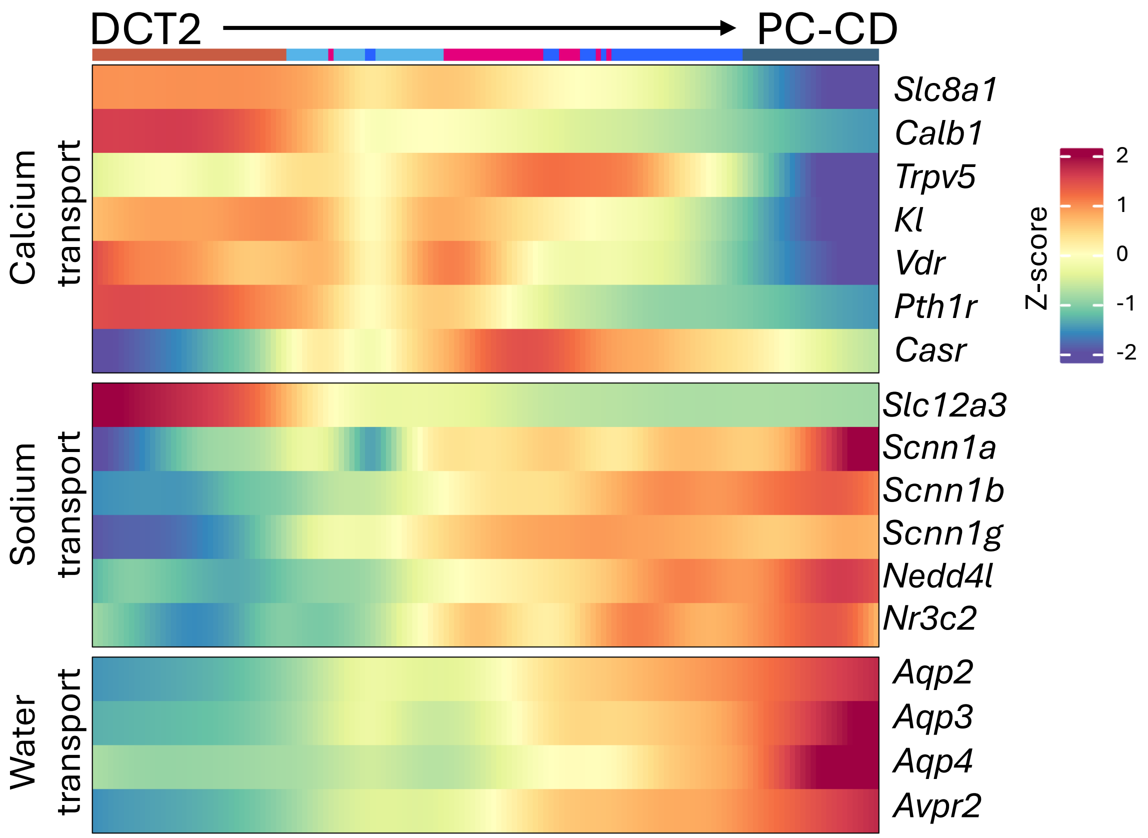

Supplemental Figure 16

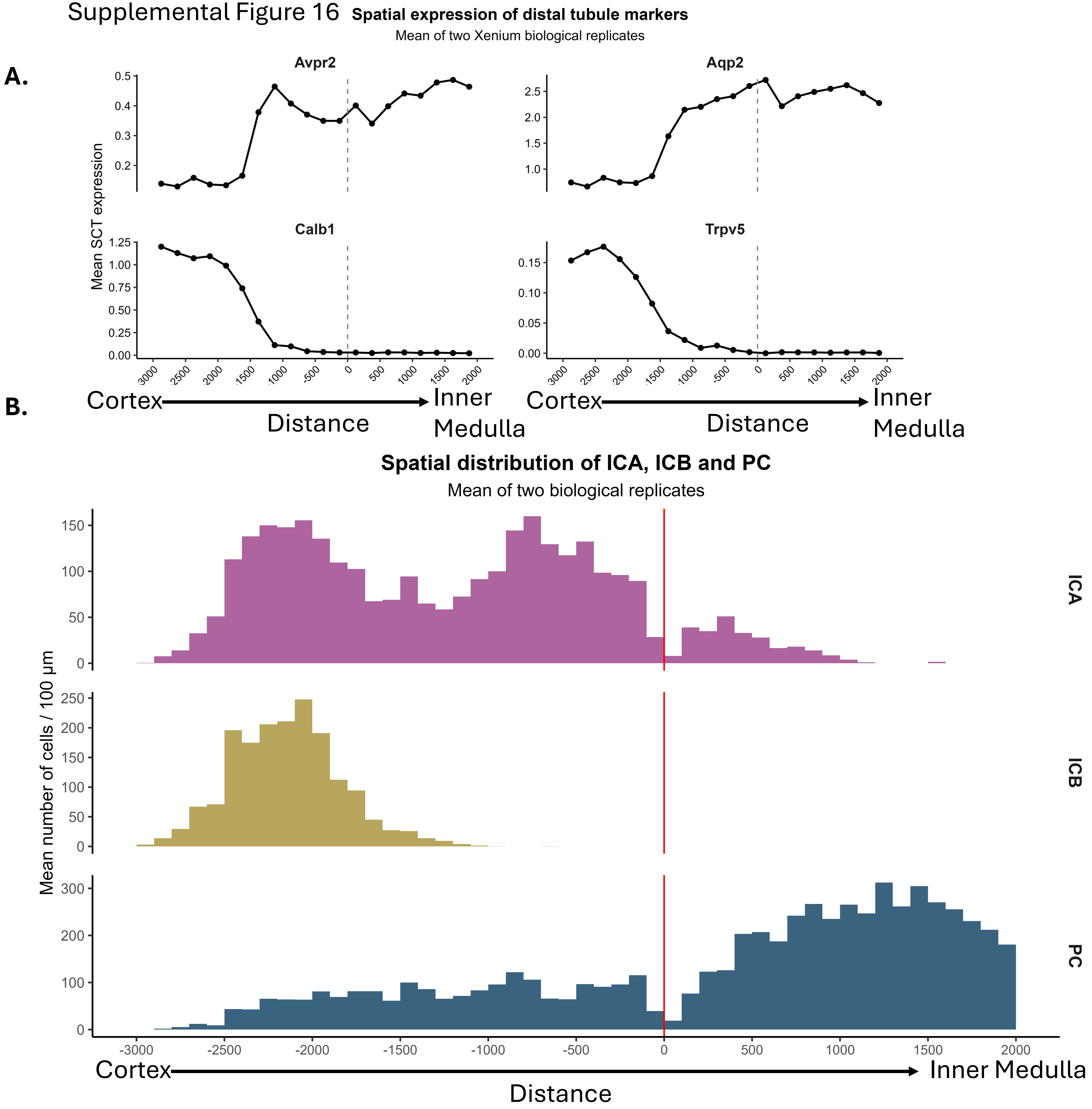

### Supplemental Figure 17

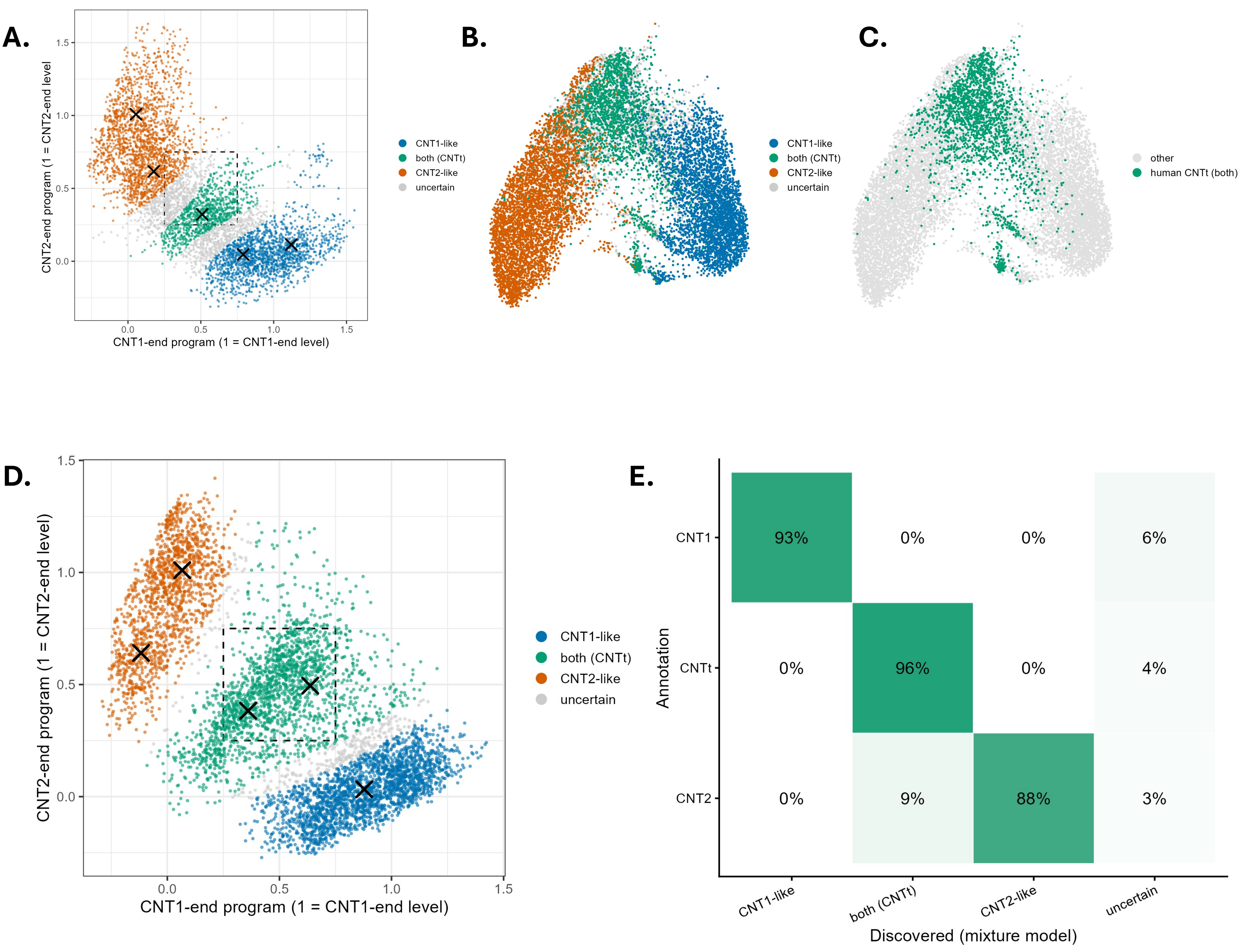

Supplemental Figure 18

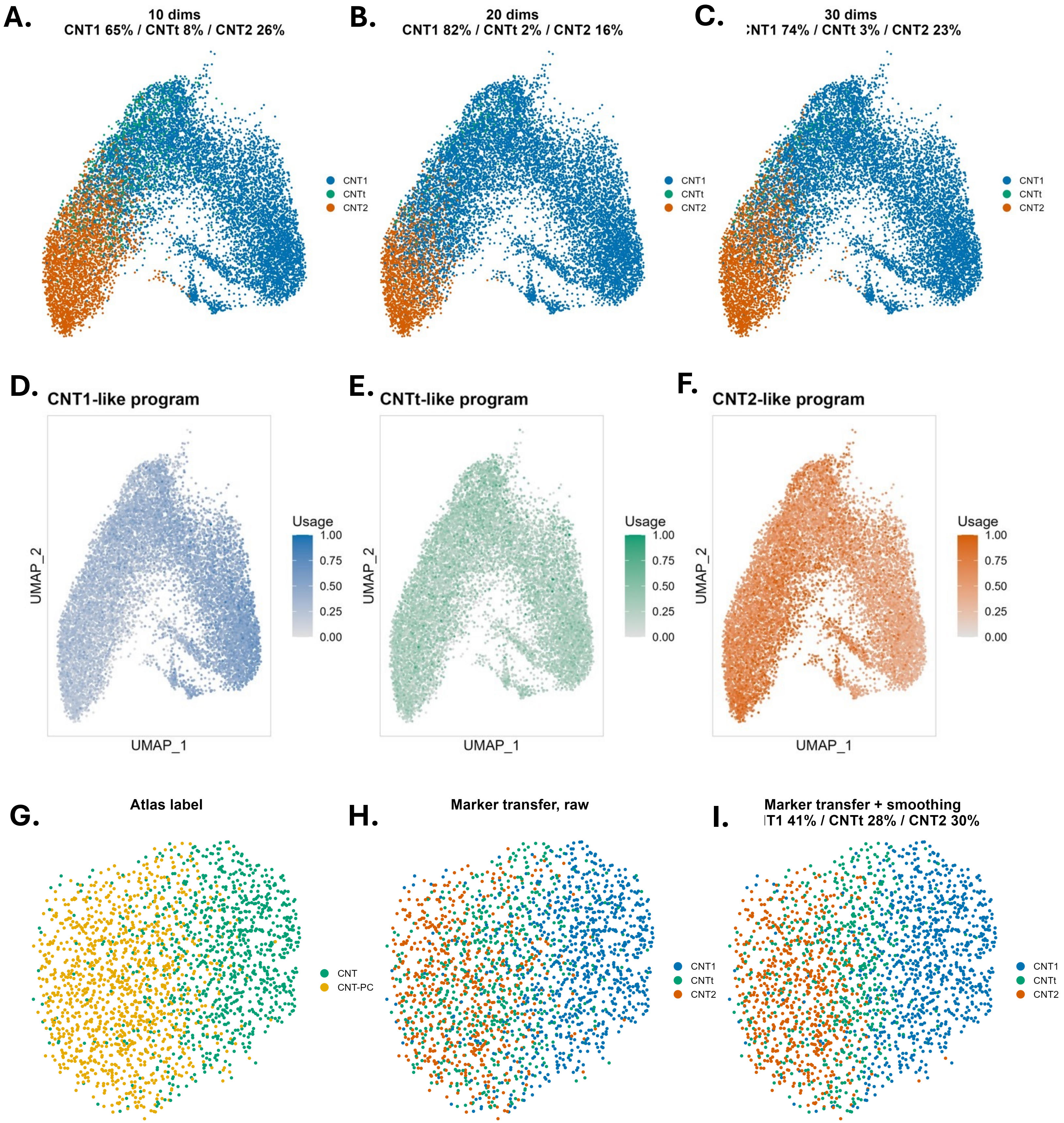

### Supplemental Figure 19

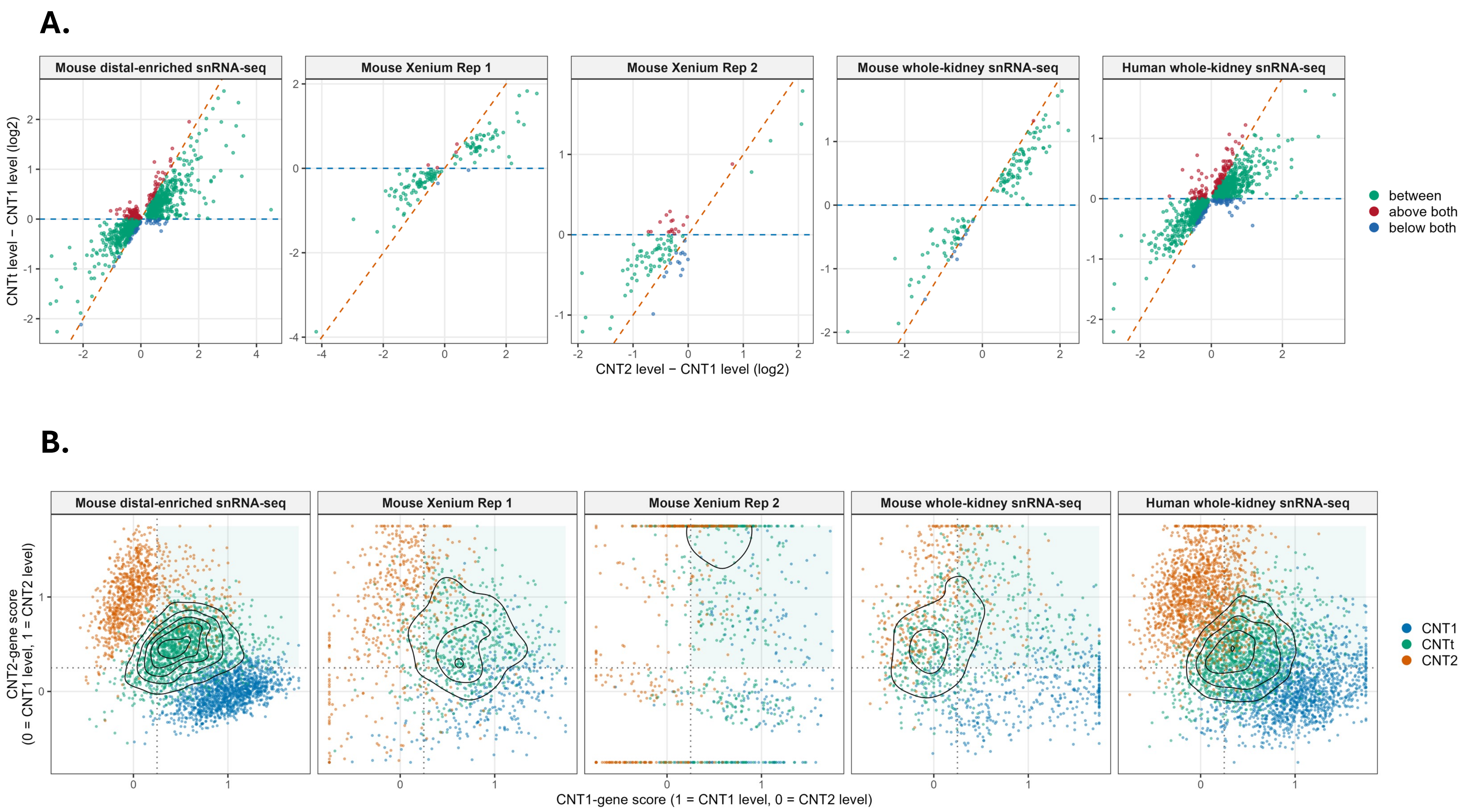
